# Evaluating agentic AI for biological discovery in autonomous and copilot settings

**DOI:** 10.64898/2026.06.04.729919

**Authors:** Shreya Johri, Erica Pimenta, Josephine Yates, Jingxin Fu, Erik L. Bao, Hyeji Jun, Brendan Reardon, Sasha Bacot, Maha Shady, Doris Fu, Wenbin Mei, Sabrina Y Camp, Jihye Park, Eliezer Van Allen

**Author notes:** co-senior authors (Jihye Park, Eliezer Van Allen).

## Abstract

Advances in large language models (LLMs)-based artificial intelligence (AI) agents have improved their ability to execute structured analytical workflows, including standard bioinformatic pipelines for biological discovery. However, computational biology rarely consists of deterministic pipeline execution alone. Biological datasets are heterogeneous and noisy, and meaningful discovery often requires open-ended hypothesis generation and iterative reasoning over multimodal evidence. These challenges are particularly evident in multi-omic studies, where paired molecular modalities and heterogeneous clinical contexts create both opportunities and obstacles for discovery. The extent to which emerging agentic AI systems can support or automate this mode of scientific discovery remains poorly understood. Here, we systematically evaluated the capabilities and limitations of agentic AI for biological discovery using multi-omic single cell datasets spanning 11 cancer types. We developed the Multistep Multimodal Multiomic Agentic (M3A) Framework to support LLM-driven reasoning over persistent multimodal data states and to capture agentic reasoning behavior in autonomous and human-AI copilot settings. Using this framework, we assessed AI agents across complementary tasks, including autonomous cell-type annotation, generation of falsifiable biological hypotheses from gene programs, and copilot experiments testing the effect of human involvement and domain expertise. We found that current AI agents are effective at broad, systemic exploration of complex data, whereas domain experts remain critical for methodological guidance and biological synthesis across analyses. Together, our results delineate the current potential and boundaries of agentic AI in computational biology, and establish a framework for evaluating AI systems designed to support biological discovery.

## Introduction

Large language model (LLM)-based AI agents have emerged as a promising paradigm for biological discovery. By integrating natural language reasoning with dynamic tool invocation, these systems can orchestrate specialized bioinformatic pipelines, query curated biological databases, and synthesize rapidly evolving scientific literature within a unified computational framework^1–6^. Complementary progress in reinforcement learning and alignment research^7,8^ has sharpened their capacity for multi-step reasoning and coordination across heterogeneous scientific workflows^3,9^, positioning them as powerful engines for biological discovery.

AI agents for biology have been evaluated across an expanding range of tasks, from knowledge retrieval activities such as literature summarization, database querying, and supplementary data extraction^10^, to bioinformatic workflows including gene-set enrichment^11^, gene-trait association^12^, gene expression preprocessing^13^, and pipeline orchestration spanning transcriptomic, epigenomic, and chromatin accessibility modalities^14,15^. In the single-cell and spatial domains (tasks that emphasize high dimensional data analysis and multi-modal learning), agents have been assessed on cell-type annotation^6,15–17^, spatial domain segmentation^18,19^, and scientific insight retrieval from published studies^20^, with synthetic data augmentation and metadata scrambling increasingly adopted to enforce objective evaluation^21^. Parallel efforts have pushed toward fully autonomous research pipelines capable of scientific software generation and single-cell method discovery.^2^

Yet, existing evaluations share several structural limitations that leave key questions about agentic biological reasoning unaddressed. Prior work focused on either single modality bioinformatic analyses typically using scRNA datasets, or pre-defined multi-omic workflow orchestration where task success reflects instruction-following capacity rather than scientific judgment. Reasoning capabilities that require iterative integration of complex data types, such as single cell multiome (scMultiome; scRNA and scATAC) datasets, remain largely unexplored. Where agents have been evaluated on discovery tasks, validation has relied on matching a single hypothesis from the originating study or qualitative expert review of a small number of case studies within the same dataset, neither of which can establish whether findings reflect genuine insights that validate across datasets. Lastly, while autonomous operation has been extensively studied, how real-time human involvement and domain expertise modulate reasoning quality in biology remains unexamined.

Here, we systematically assessed the quality of AI agents’ multi-step, multimodal, and multi-omic reasoning in real-world biological tasks. As a testbed, we utilized cancer scMultiome data (11 cancer types, 182 patients, 1,064,365 cells) from 4 published studies to enable and explore reasoning over paired RNA and ATAC modalities spanning patient cohorts with complex disease biology and heterogeneous clinical contexts. To enable these systematic evaluations, we developed the Multistep Multimodal Multiomic Agentic (M3A) Framework, a standardized environment for LLM-driven agentic reasoning that maintains persistent multimodal data states and captures step-level telemetry during autopilot and copilot modes. With these multi-modal cohorts and the M3A framework, we assessed autonomous cell-type annotation for scMultiome datasets, tested whether agent-generated biological hypotheses regarding cell programs (treated as falsifiable hypotheses) generalized to independent cohorts across 64 tasks, and examined how human involvement and domain knowledge alter agentic behavior and performance through human-AI copilot experiments.

## Results

We developed an agentic framework to measure how LLMs perform data-driven reasoning over complex computational biology workflows (Fig. 1; Methods). The Multistep Multimodal Multiomic Agentic (M3A) framework enables controlled execution of LLM agents while capturing their full analytical trajectories, providing a foundation for both quantitative performance evaluation and mechanistic analysis of reasoning. It enforces constraints across five core dimensions within a unified system, specifically designed to enable rigorous tracing of agent behavior to match the diverse demands of complex biological workflows: (i) a standardized execution environment that ensures reproducible baseline conditions; (ii) a unified tool suite spanning literature retrieval, programmatic analysis, terminal access, and domain-specific single-cell pipelines; (iii) recursive multimodal context integration that incorporates model outputs across steps; (iv) persistent data state that maintains structured objects across tool invocations; and (v) step-level telemetry that captures intent, tool selection, and outcomes at each decision point (Methods). Together, these components ensure that model behavior is reproducible, directly comparable across runs, and amenable to mechanistic analysis.

**Figure 1:**
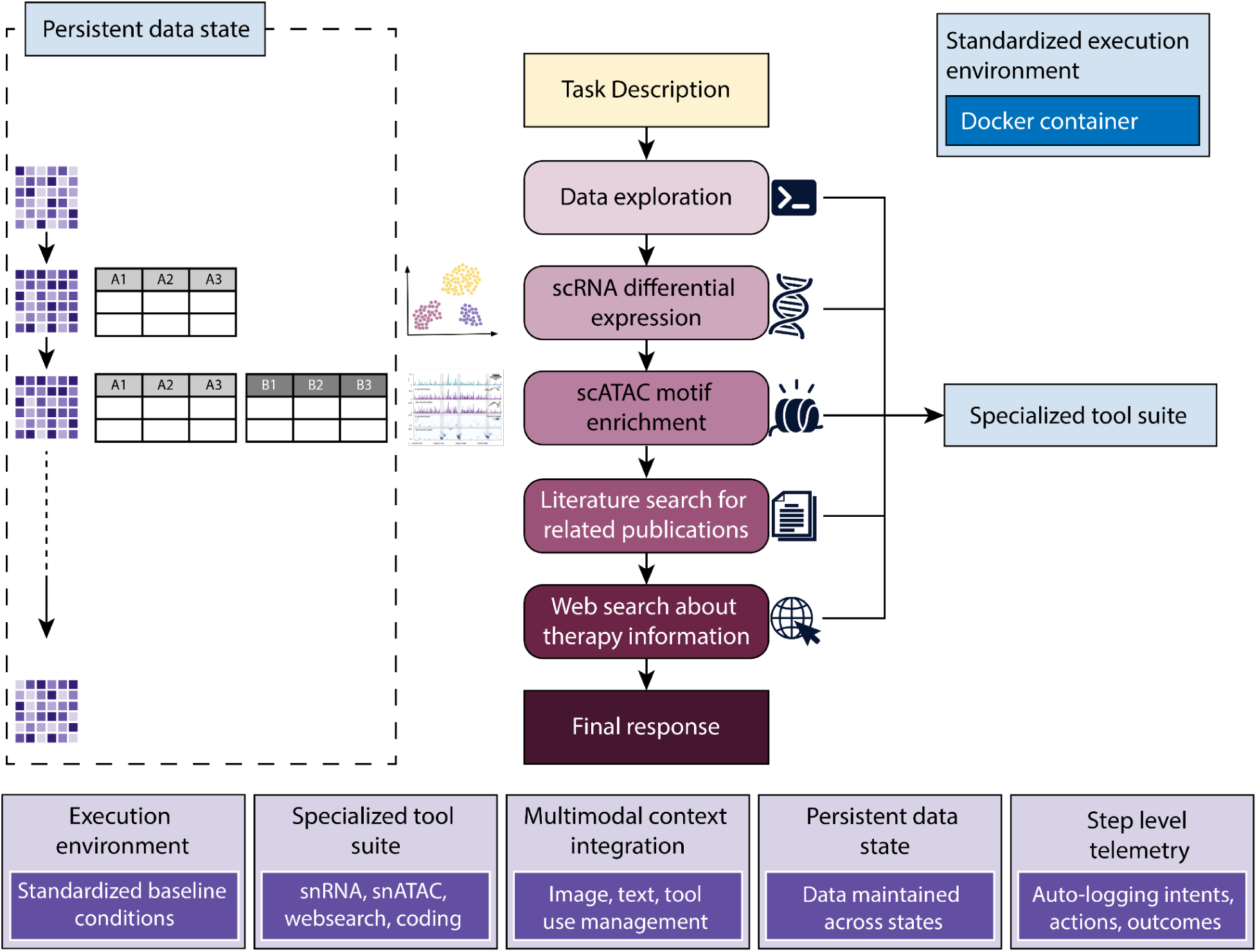
The Multistep Multimodal Multiomic Agentic (M3A) evaluation framework.

We applied the M3A framework to systematically evaluate a Claude Opus 4.6^22^ (version Feb 6, 2026) based AI agent (M3A agent) across multi-step, multi-omic reasoning tasks with predefined evaluation endpoints spanning molecular profiling data from multiple cancer types (Fig. 1, Supplementary Table 1). We assessed four core questions: how well the agent characterizes complex datasets through multi-step reasoning and data analysis; how robust its findings are when applied to external datasets; how human intervention shapes reasoning dynamics; and what level of domain expertise benefits most from AI assistance for discovery tasks.

### AI agent-driven cell type characterization is limited for rare cell-types

We first evaluated whether an autonomously operating M3A agent can perform a complete cell type annotation workflow using multi-patient single-nucleus multiome (RNA expression and ATAC profiling; snMultiome) disease datasets. To this end, we selected 146 samples from 130 patients across 9 cancer types from the Human Tumor Atlas Network (HTAN)^23^ pan-cancer snMultiome dataset (n = 646,057 cells). These data included paired snMultiome profiles as well as matched modality-specific libraries, including snRNA-seq and snATAC-seq (Supplementary Table 1). Ground-truth (GT label) cell type annotations were previously established by consensus from individual HTAN cancer-type working groups.^24,25^

We evaluated the M3A agent’s cell type annotations at two levels of resolution: (i) broad cell-type classes and (ii) fine-grained cell-type labels within a given class. For broad-level evaluation, each author-provided GT label and M3A agent generated label (AI label) were mapped to six major categories: epithelial, stromal, endothelial, immune, neural and cycling (Supplementary Note 1). For fine-grained evaluation, cancer-type-specific hierarchical ontologies were constructed from GT labels, and each AI label was manually mapped to a node in the corresponding ontology. Calls were then scored by comparing the position of the AI label to the GT label within the ontology, yielding six mutually exclusive tiers ranging from exact match to multiple types of annotation errors (Methods).

For the broad cell-type annotation task, the M3A agent achieved high specificity (mean = 0.975) but comparatively lower sensitivity (mean = 0.853) across different cancers (Fig. 2a, Supplementary Table 2, Supplementary Figure 1; Methods). Specificity was consistently high across cancer types, with the highest observed in Uterine Corpus Endometrial Carcinoma (UCEC) (=0.991) and the lowest in Glioblastoma (GBM) (=0.938), indicating few false-positive annotations. In contrast, sensitivity varied more substantially across datasets, ranging from highest in UCEC (=0.974) to lowest in GBM (=0.330), suggesting that recall is more dependent on disease context and dataset complexity.

**Figure 2:**
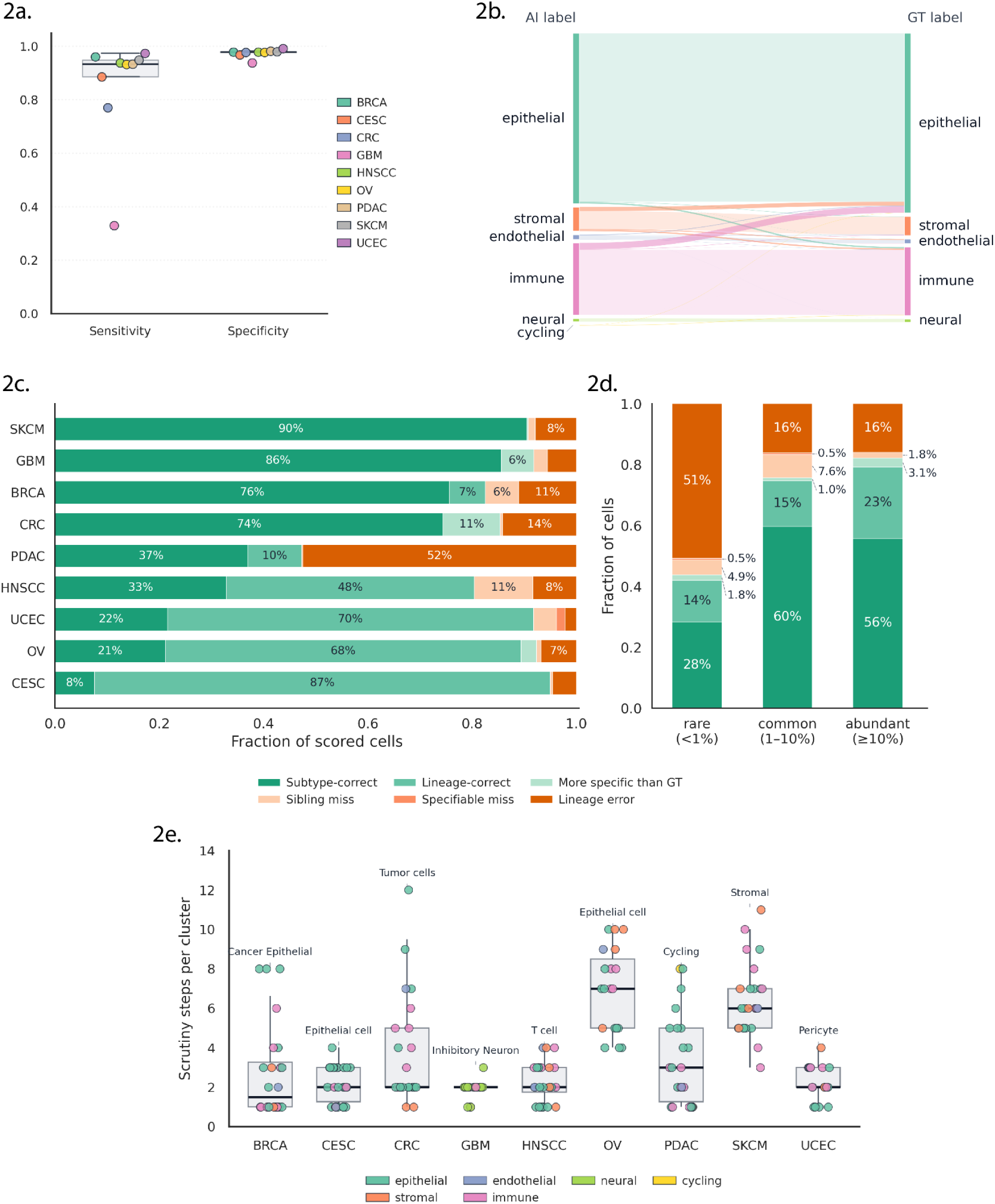
Pan-cancer evaluation of M3A agent’s cell-type annotations against author annotations across nine cancer-types: Breast Cancer (BRCA), Cervical Squamous Cell Carcinoma (CESC), Colorectal Cancer (CRC), Glioblastoma (GBM), Head and Neck Squamous Cell Carcinoma (HNSCC), Ovarian Cancer (OV), Pancreatic Ductal Adenocarcinoma (PDAC), Skin Cutaneous Melanoma (SKCM), Uterine Corpus Endometrial Carcinoma (UCEC). **(a)** Macro-averaged sensitivity and specificity at the broad cell-type tier (epithelial, stromal, endothelial, immune, neural, cycling), colored by cancer-type. For each cancer-type, sensitivity and specificity were computed for every broad class by one-vs-rest decomposition of the multi-class confusion matrix, then averaged uniformly across classes present in the cohort box center line, median; box bounds, 25th-75th percentiles (IQR); whiskers, most extreme cohort within 1.5× IQR. **(b)** Pan-cancer Sankey diagram of M3A agent’s predictions (left blocks) flowing to ground truth (GT) annotations (right blocks) at the broad cell-type tier. Block height is proportional to the number of cells in each broad category. **(c)** Per-cohort decomposition of scored cells into six scoring conditions: Subtype-correct, Lineage-correct, More specific than GT, Sibling miss, Specifiable miss, and Lineage error (Methods). **(d)** Scoring condition composition stratified by class prevalence: rare <1%, common 1-10%, abundant ≥10%. Each bar shows the fraction of cells in six mutually exclusive conditions: Subtype-correct, Lineage-correct, More specific than GT, Sibling miss, Specifiable miss, and Lineage error. **(e)** Per cancer-type distribution of M3A agent scrutiny depth across single-cell clusters. Each cohort’s boxplot summarizes the number of reasoning iterations (“scrutiny steps”) the M3A agent took before committing to a cell-type label, computed independently for every cluster the M3A processed in that cohort. Each dot is one cluster, colored by the broad cell-type category of the M3A assigned label. Box center line, median; box bounds, 25th-75th percentiles (IQR); whiskers, most extreme cluster within 1.5×IQR. The cluster with the highest scrutiny depth in each cohort is annotated with its M3A assigned label.

Overall, the AI label agreed with the GT label for 94.0% of scored cells (Fig. 2b, Supplementary Table 2). Misclassifications were non-randomly distributed: errors were directional rather than symmetric, with epithelial-GT cells being assigned immune or stromal AI labels (2.3% and 1.5%) far more often than the reverse (immune-GT cells assigned an epithelial label at 0.5%, and stromal-GT cells at 0.1%). A further subset of discordant assignments converged on the stromal AI label, with immune-GT and endothelial-GT cells assigned a stromal AI label in 0.5% and 0.3% of all scored cells, respectively. Smaller discrepancies included epithelial-GT cells assigned an endothelial AI label (0.2%) and stromal-GT cells assigned an immune AI label (0.1%), with the remaining 0.4% distributed across other minor cross-type assignments.

For the fine cell-type level annotation tasks, accuracy varied substantially across cancers, with much of the accuracy loss attributable to vocabulary gaps rather than outright misclassification (Fig. 2c, Supplementary Table 2; Methods). Scoring was assessed using six mutually exclusive conditions: subtype-correct (exact match), more specific than GT, lineage-correct, lineage error, specifiable miss, and sibling miss. The fraction of subtype-correct cells ranged from 7.6% in CESC to 90.5% in SKCM (mean = 49.6%). For the four lower-accuracy cohorts (CESC, UCEC, OV, HNSCC), the predominant non-exact tier was lineage-correct but vocabulary-limited (87.5%, 70.2%, 68.2%, and 47.6% of cells, respectively), indicating that the M3A agent placed cells on the correct lineage but did not have a finer term available in its working vocabulary. Lineage errors were rare across most cancer types but markedly elevated in PDAC, where 52.5% of cells received a label outside the GT lineage. Specifiable misses and sibling misses together accounted for fewer than 12% of cells in every cohort, suggesting that when the M3A agent did possess the fine-grained vocabulary, it generally applied it correctly.

Per-class fine accuracy depended strongly on class prevalence, with rare cell types substantially harder to annotate than common or abundant ones (Fig. 2d, Supplementary Table 2). We stratified cells by the prevalence of their GT label within each cohort: rare (<1%), common (1-10%), and abundant (≥10%). Correct calls (subtype-correct, more specific than GT, or lineage-correct combined) accounted for only 43.8% of rare cells but rose to 75.7% for common and 82.2% for abundant classes. The dominant failure mode for rare classes was cross-family error: 50.8% of rare cells received a label outside the GT lineage, compared with 16.2% and 16.0% for common and abundant classes.

We next investigated whether the number of analytical steps the M3A agent took per cluster for cell type assignment translated into higher annotation accuracy (Fig. 2e, Supplementary Table 2). Per-cluster scrutiny ranged from 1 to 12 steps, with per-cohort medians spanning 1.5 (BRCA) to 7.0 (OV). Two cohorts received consistently deeper scrutiny by the M3A agent than the rest: OV (median = 7.0, IQR 5.0-8.5) and SKCM (median = 6.0, IQR 5.0-7.0), while the remaining seven cohorts had median values between 1.5 and 3.0. Within-cohort variability was largest in CRC (range 1-12) and SKCM (range 3-11). Across clusters, deeper M3A agent scrutiny was not associated with higher annotation accuracy (Supplementary Figure 1). Scrutiny showed no relationship with either broad-tier accuracy or fine-tier accuracy, suggesting that scrutiny reflects label ambiguity rather than a driver of correctness (see Methods).

Finally, qualitative analysis of reasoning traces revealed that although scATAC was universally used, M3A agent heavily steered towards using it for label confirmation rather than independent clustering. Additionally, marker-gene annotation drew solely on model knowledge without web retrieval, quality control steps were never revisited, and iteration was limited to refining cluster identities (Supplementary Figure 2). These findings indicate that without explicit procedural instructions, M3A agents often defaulted to one-shot analysis rather than the iterative workflows often employed by human computational biology scientists.

### Validation tasks reveal limitations in AI-agent driven prioritization of biological programs

Evaluating discovery is fundamentally more challenging than evaluating bioinformatic workflows with well-defined outputs, because ground-truth labels are inherently incomplete for discovery. Any published paper captures only one of many possible discoveries from a dataset (hence only one ground-truth label), and expert consensus (even if feasible) is costly, time-consuming, and prone to reflect prior expectations rather than empirical evidence. We therefore designed a set of tasks that evaluate reasoning indirectly, treating agent-generated hypotheses as falsifiable predictions and assessing their concordance with held-out data (Fig. 3a; Methods). This approach is scalable and objective since ground-truth labels are computed automatically from independent data, requiring no human annotation, and concordance with held-out data ensures findings are robust rather than dataset-specific.

**Figure 3:**
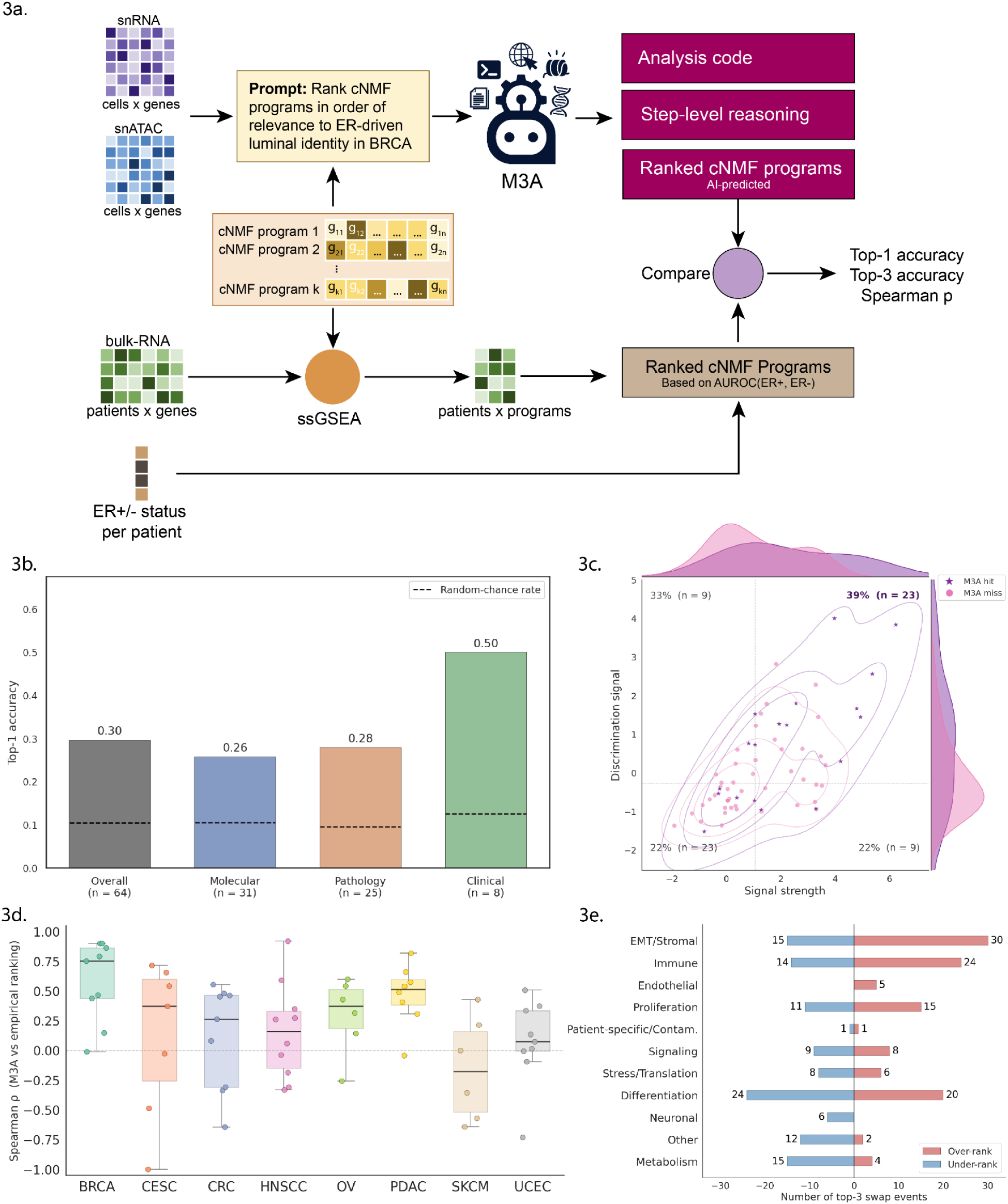
Concordance between M3A agent program rankings and statistical ground-truth rankings. **(a)** Schematic representing an example task in BRCA, illustrating the benchmark design applied across n = 64 tasks in 8 TCGA cancer types (BRCA, CESC, CRC, HNSCC, OV, PDAC, SKCM, UCEC). **(b)** Top-1 accuracy (fraction of tasks for which M3A ranked the statistically top-scoring program first) across all 64 tasks, stratified by endpoint category. Dashed lines indicate the task-weighted random-chance baseline for each category. **(c)** Top-1 accuracy as a function of task structure. Each point represents one task; x-axis, signal strength (within-metric-type percentile rank of the best program’s validation score); y-axis, discrimination signal (score gap between the first- and second-ranked programs). Quadrant labels show top-1 accuracy and task count. **(d)** Spearman rank correlation between M3A agent and empirical program rankings per cancer type. Each point represents one task; box, interquartile range; horizontal line, median; whiskers, 1.5× IQR. Dashed line at ρ = 0. **(e)** Directional ranking bias by biological program theme. Bars show the number of top-3 swap events in which a program of that theme was promoted into the M3A agent’s top-3 but absent from the empirical top-3 (over-rank) or present in the empirical top-3 but absent from the M3A agent’s top-3 (under-rank). Numbers indicate swap event counts.

Specifically, consensus non-negative matrix factorization (cNMF)^26^ was performed on HTAN snRNA tumor compartment data, and the resulting unannotated gene programs were provided to M3A agent alongside paired snRNA and snATAC. The M3A agent was prompted to rank the programs by relevance to a defined endpoint purely through reasoning, thereby producing falsifiable hypotheses about which programs should associate with independent molecular, pathological, or clinical phenotypes. The same cNMF programs were scored in an independent TCGA bulk RNA-seq cohort via ssGSEA and ranked using endpoint-appropriate metrics (AUROC for binary molecular and pathological endpoints, Spearman correlation for continuous measures, and Cox proportional hazards for survival) providing ground-truth rankings derived from held-out empirical data rather than human judgment. Concordance between the M3A agent’s ranking and the empirically-derived ranking measured biological reasoning accuracy, while the spread of scores (based on endpoint-appropriate metrics) across programs quantified task difficulty. Tasks were selected based on available endpoints across six modalities orthogonal to transcriptomics, representing cancer-type specific biology: DNA mutations, copy number alterations, DNA methylation, immunohistochemistry (IHC) status, histopathology status, and clinical survival. A significant positive rank correlation indicated that the agent’s prioritization, derived from snMultiome data, generalized to an independent bulk RNA-seq data from TCGA (Fig 3a, Supplementary Table 3; Methods).

Across 64 benchmark tasks, M3A agent identified the independently assessed top-ranked cNMF program as its first ranked prediction (top-1 accuracy) in 30% of cases, nearly three-fold above the task-weighted random-chance rate of ∼11% (Fig. 3b, Supplementary Table 3). This was consistent across endpoint categories: top-1 accuracy was 26% for molecular endpoints (n = 31), 28% for pathology (n = 25), and 50% for clinical/survival endpoints (n = 8), with all categories substantially exceeding their respective baselines. Top-1 accuracy was strongly driven by the nature of the empirical signal (Fig. 3c, Supplementary Figure 3, Supplementary Table 3). When both signal strength (the percentile rank of the best program’s validation score within its metric type) and discrimination (the score gap between the first- and second-ranked programs) were simultaneously high, top-1 accuracy reached 39% (n = 23 tasks); when either dimension was low, accuracy fell substantially (low signal, low discrimination: 22%, n = 23; low signal, high discrimination: 33%, n = 9; high signal, low discrimination: 22%, n = 9). Example tasks are shown in Supplementary Figure 3.

Overall ranking performance, measured by Spearman correlation, varied substantially across cancer types (Fig. 3d, Supplementary Table 3). BRCA exhibited the highest median Spearman rank correlation between M3A agent and empirical rankings (ρ = 0.75), followed by PDAC (ρ = 0.50) and CESC and OV (ρ = 0.38), all cancer types with well-characterised transcriptional programs and strong prior literature coverage. However, ranking performance for SKCM showed near-zero to negative median correlations (ρ = −0.2). Analysis of ranking errors by biological program theme, measured by top-3 cNMF program swap events, revealed that failures were structured and interpretable rather than random (Fig. 3e, Supplementary Table 3). EMT/Stromal and Immune programs were the most systematically over-ranked by the agent (30 and 24 over-rank events, respectively). Conversely, Metabolism and Neuronal programs were consistently under-ranked (15 and 6 under-rank events, respectively), pointing to thematic blind spots in the agent’s encoded biological priors. Differentiation and Signaling programs showed near-symmetric over- and under-ranking (Differentiation: 20 over, 24 under; Signaling: 8 over, 9 under), indicating variable but unbiased treatment. Together, these results establish that M3A agent’s reasoning errors trace to specific, interpretable gaps in prior knowledge rather than arbitrary failures.

### Copilot experiments reveal distinct behavioral profiles and shifted analytical priorities in M3A

Given our benchmarks revealed the varying reasoning strengths and limitations of AI-agents with real-world cancer biology tasks, we subsequently designed a human-AI copilot experiment to identify whether and how components of computational biology workflows could be automated in the context of varying human technical expertise and domain knowledge (Fig. 4a, Supplementary Table 4; Methods). We first established three interaction modes for M3A agents that represent a decreasing level of human engagement with the agents: Human-in-the-Loop (HITL), Human-on-the-Loop (HOTL), and fully autonomous. We then applied these strategies to three additional clinically integrated multi-omics cancer biology datasets: liposarcoma (LPS)^27^, esophageal adenocarcinoma (EAC)^28^, and breast cancer (BRCA)^29^. A representative cohort of human computational biologists was stratified into three experience groups for HITL and HOTL runs: (i) dataset experts (the first authors of the LPS, EAC, and BRCA studies from which the datasets were drawn); (ii) single-cell experienced analysts (researchers with one or more years of active single-cell RNA-seq project experience); and (iii) single-cell novices (researchers with no single-cell experience or fewer than six months of exposure). Through HITL or HOTL setups, they (as well as the fully autonomous agent) performed four biological reasoning tasks of increasing complexity using the three datasets: (i) cNMF program annotation, (ii) hypothesis formulation, (iii) regulatory driver analysis, and (iv) biology synthesis. Of note, our copilot experiment was structured around extensive expert-guided curation, so even fully autonomous runs inherently leveraged human domain expertise embedded within the experimental architecture (Methods).

**Figure 4:**
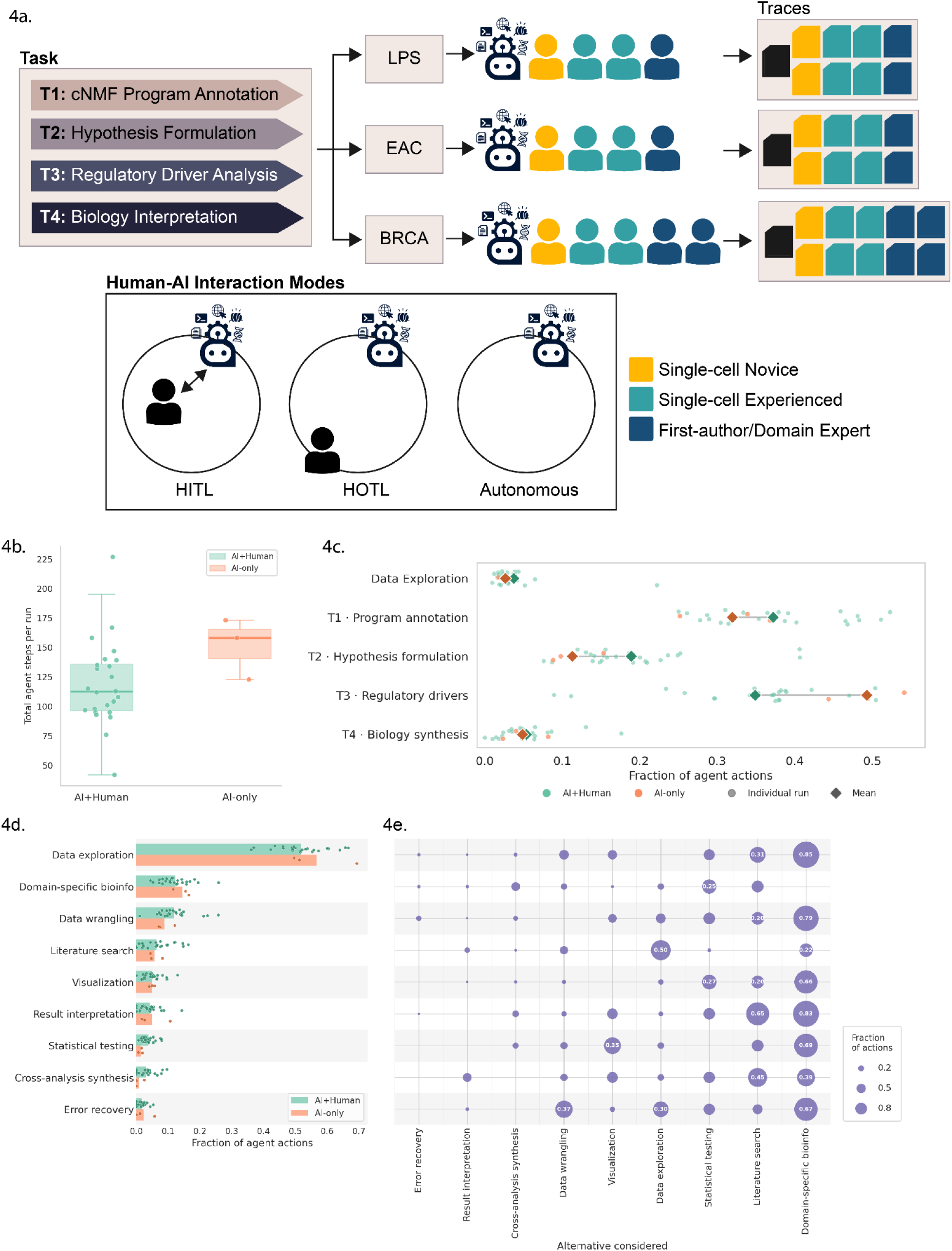
User behavior in copilot experiments. **(a)** Schematic showing the copilot experiment setup. Three interaction modes: Human-in-the-Loop (HITL), Human-on-the-Loop (HOTL), Fully autonomous, representing a decreasing level of human engagement with the agents. Three datasets:liposarcoma (LPS), esophageal adenocarcinoma (EAC), breast cancer (BRCA) datasets; outputs and traces collected. Three experience groups: dataset experts, single-cell experienced analysts, and single-cell novices. Four biological reasoning tasks of increasing complexity: cNMF program annotation, hypothesis formulation, regulatory driver analysis, and biology synthesis. **(b)** Total agent steps per run. Box shows median and interquartile range; whiskers extend to 1.5 × IQR; individual runs are overlaid as jittered points coloured by interaction mode (AI+Human or AI-only). **(c)** Distribution of M3A agent actions across sub-tasks for AI+Human (HOTL and HITL combined) and AI-only conditions. Each circle represents one run, diamond markers show group means. The x-axis gives the fraction of all agent actions in a given run attributed to each sub-task (Dataset exploration; T1, program annotation; T2, hypothesis formulation; T3, regulatory drivers; T4, biology synthesis). **(d)** Horizontal box plots showing the fraction of agent actions spread across agent action categories. Two groups per row correspond to AI+Human and AI-only. Box shows median and interquartile range; whiskers extend to 1.5 × IQR; individual runs are overlaid as jittered points coloured by interaction mode. **(e)** Alternative actions considered by the M3A agent (x-axis) for M3A agent action (y-axis). For each action category executed (rows, ordered as in left panel), bubble area represents the fraction of that action type’s executions in which the agent’s reasoning log explicitly named the column category as an alternative it considered but did not take. Bubble area is proportional to fraction; values ≥ 0.18 are annotated.

We first characterized a broad baseline of how the M3A agent distributed its analytical effort across the workflow by comparing AI+Human sessions (HOTL and HITL combined; n=24 runs) to fully autonomous AI-only runs (n=3). Overall, fully autonomous runs were longer than human-supervised sessions (median 151.3 vs. 118.5 steps, respectively; Fig. 4b; Supplementary Table 4). In AI+Human sessions, M3A agent effort was concentrated in the early analytical stages: program annotation and hypothesis formulation accounted for a mean of 37.2% and 18.9% of all agent actions per run, respectively, with regulatory driver analysis comprising an additional 34.9% of all agent actions per run (Fig. 4c, Supplementary Table 4). Fully autonomous runs showed a markedly different allocation, devoting a greater fraction of their actions to regulatory analysis (49.3% vs. 34.9%) at the expense of program annotation (11.3% vs. 18.9%). These results suggest that without human guidance, agents compress the iterative annotation step that grounds downstream biological reasoning.

To further characterize the technical breadth of each session, we classified all M3A agent actions into functional categories that span data preparation workflows (data exploration, data wrangling), specialized biological computation (bioinformatics, literature search), domain-agnostic analysis (statistical testing, visualization), knowledge-driven evaluation (single result interpretation, cross-analysis synthesis), and code troubleshooting (error recovery) (Fig. 4d, Supplementary Table 4; Methods). Data exploration dominated in both human-supervised and autonomous conditions, and it accounted for a comparable fraction of agent actions across modes (56.7% in AI-only runs vs. 51.9% in AI+Human; Fig. 4d, Supplementary Table 4). The near-complete absence of cross-analysis synthesis steps in fully autonomous runs (0.8% in AI-only runs vs. 3.1% in AI+Human) indicated that autonomous agents rarely performed integrative reasoning that synthesizes findings across sub-analyses, a task that human collaborators actively prompted. Domain-specific bioinformatics analyses were correspondingly more frequent in autonomous runs (14.5% in AI-only runs vs. 12.2% in AI+Human), potentially reflecting compensatory technical depth in the absence of directed biological interpretation. Decomposing these differences by task revealed the largest divergences were observed in data exploration followed by domain-specific bioinformatics in the regulatory driver analysis task (Supplementary Figure 4).

To investigate how the M3A agents reason through multi-step processes, we next analyzed their consideration of alternative actions that were not ultimately pursued (Fig. 4e, Supplementary Table 4). Across all M3A agent actions, domain-specific bioinformatics tool usage was considered as an alternative action in 85.1% of data exploration actions, 83.0% of result interpretation actions, and 79.0% of data wrangling actions, indicating that a pivot toward domain-specific analysis was a frequently contemplated alternative path at most steps in the workflow but rarely executed. Literature search was the second most commonly considered alternative action, appearing alongside 65.2% of result interpretation actions and 44.7% of cross-analysis synthesis steps. Notably, error recovery (<3%), result interpretation (<10%), and cross-analysis synthesis (<10%) were rarely considered as alternatives.

Co-occurrence patterns revealed that M3A agents rarely considered a single alternative reasoning pathway; rather, alternative reasoning strategies clustered around domain-specific bioinformatics as an anchor (78.3% of all agent actions; Supplementary Figure 4). Other alternatives, like literature search (30.4%), data exploration (21.0%), and statistical testing (18.4%), were also frequently considered. Interestingly, error recovery (2%), result interpretation (1%), and cross-analysis synthesis (4%) were rarely considered. The most frequent jointly considered pairs of alternative reasoning strategies all featured a domain-specific bioinformatics anchor paired with literature search (22.9%), data exploration (13.6%), and statistical testing (12.1%), forming a core triad at nearly every analytical branch point. This alternative prioritization suggests that the AI agents’ decision-making framework is built upon a domain-specific bioinformatics foundation, pointing to their technical emphasis over biological interpretation.

### Interaction mode and user experience influence human steering strategies and agentic focus

We next examined where in the agentic workflow users chose to intervene, and whether these actions depended on biological expertise (Fig. 5a, Supplementary Table 5, Supplementary Note 4-5). User-steering events were defined as direct human interventions logged during the session, comprising feedback and instructions issued to the agent, as well as session documentation. Across all experience groups, user steering was concentrated in program annotation and hypothesis formulation. Domain experts (n = 4) showed the most pronounced redistribution between interaction modes, increasing their steering during hypothesis formulation by 24.5% in HITL relative to HOTL, while reducing it at the regulatory driver analysis stage by 37.4% in HITL relative to HOTL. This behavior indicated a pattern whereby domain experts strategically front-loaded their guidance when biological reasoning was deemed most consequential and ceded the regulatory driven analysis step to the agent once the analytical direction had been established. Single-cell experienced analysts (n=13) showed stable steering profiles across modes, whereas single-cell novices (n=5) disproportionately increased their engagement at hypothesis formulation in HITL (+28.2%).

**Figure 5:**
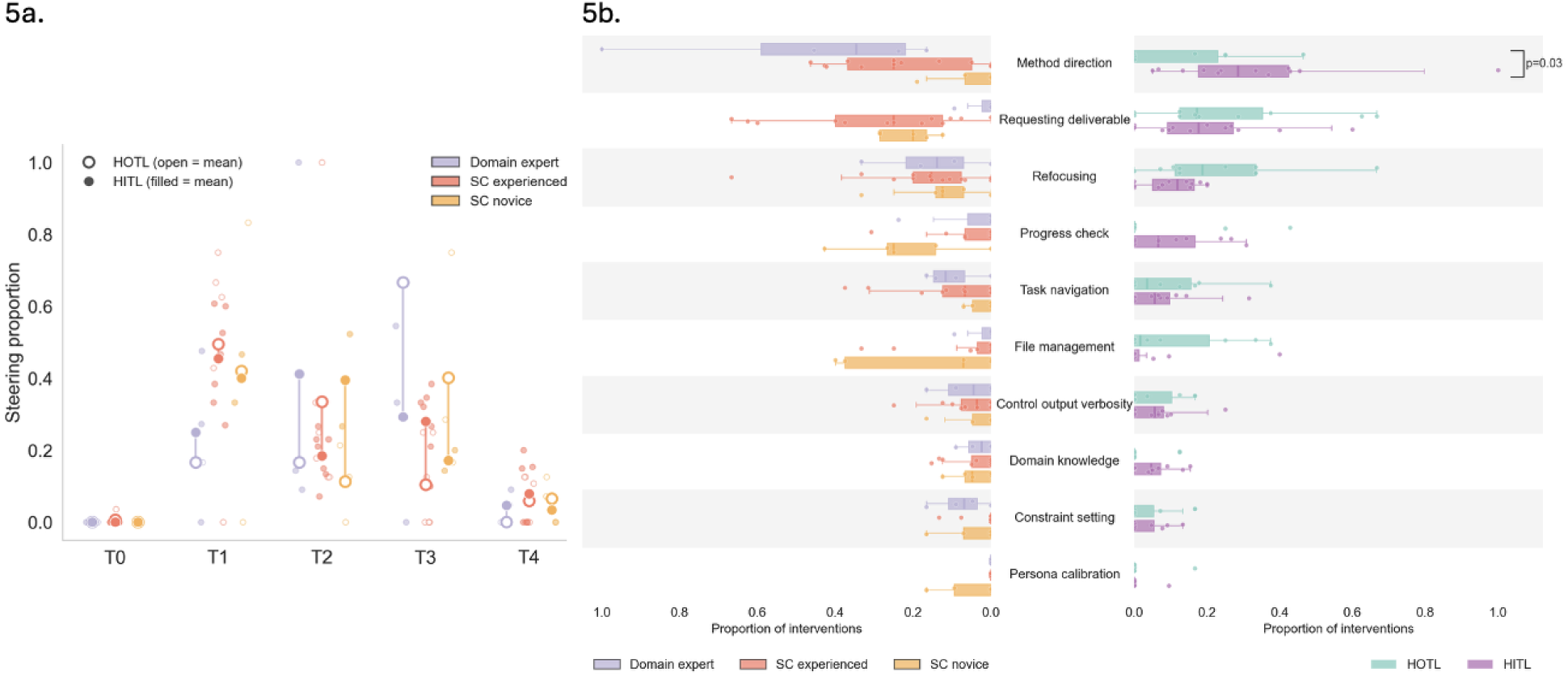
Effect of interaction mode and user experience on steering of M3A agents. **(a)** Steering proportion per task (T0; data exploration, T1; cNMF program annotation, T2; hypothesis formulation, T3; regulatory driver analysis, and T4; biology synthesis), broken down by experience level and interaction mode. Each task shows three experience groups (domain expert, SC experienced, SC novice; colour-coded) with a vertical dumbbell per group connecting the mean HOTL proportion (open circle) to the mean HITL proportion (filled circle). Translucent dots show individual run values for HOTL (open) and HITL (filled). **(b)** Human intervention patterns by experience level and interaction mode. Butterfly layout with human intervention categories on the shared central y-axis, sorted by overall mean proportion (highest at top). *(Left panel)* Horizontal box plots extending left, showing the fraction of interventions per category for each experience group (domain expert, SC experienced, SC novice). *(Right panel)* Horizontal box plots extending right, comparing HOTL and HITL intervention fractions per category. Boxes show IQR and median; jittered dots represent individual runs. Horizontal brackets indicate categories with a significant HOTL vs. HITL difference by Wilcoxon signed-rank test (paired by user × dataset; p < 0.05, uncorrected).

To further characterize human-AI interaction, user actions were annotated into categories reflecting different modes of intervention (Methods). Distinct behavioral patterns were observed when stratifying users by expertise (Fig. 5b, Supplementary Table 5): domain experts concentrated nearly half of their intervention on method direction (46.5%), whereas single-cell novices dedicated substantial effort to file management (16.9%) and progress checks (21.8%). Notably, novice users also leveraged persona calibration (5.2%) to prompt the agent for more detailed explanations, reflecting a strategic adaptation to their unfamiliarity with the datasets and downstream analyses. These expertise-dependent differences were evident not only at the aggregate task level, but also across individual subtasks (Supplementary Figure 5). Furthermore, the specific nature of human supervision dictated distinct operational profiles: closer human engagement in HITL sessions pointed agent effort toward method direction (25.1%), progress checks (9.5%) and domain knowledge (6%). In contrast, for the HOTL mode, humans pivoted the AI toward workflow refocusing (25.3%) and file management (11%) (Fig. 5b, Supplementary Table 5). These shifts suggest that real-time human engagement redirected the agent away from workflow maintenance and toward biological reasoning.

### Blinded expert reviews reveal the complementary strengths of agents and domain experts

Finally, to further explore the strengths and weaknesses of these varying agentic AI strategies for biological discovery, the originating study first authors or dataset experts conducted a blinded review of the final outputs from copilot (HITL and HOTL) and fully autonomous runs without labels indicating the interaction mode or their technical background. These reviewers scored the task quality based on scientific accuracy, depth, and evidence, provided credibility and novelty scores based on final outputs, ranked the overall outputs, and benchmarked the outputs against their own findings (Supplementary Table 6). Of note, they also attempted to infer the interaction mode and analyst expertise of each run (Methods), but their predictions were near random, indicating successful blinding (Supplementary Figure 6).

When considering all tasks, the fully autonomous run achieved the highest median performance ranking by the blinded reviewers, followed by HOTL and HITL (Fig. 6a). When evaluated by user expertise, computational analysts without single cell experience (SC Novice) and fully autonomous runs consistently clustered near the top of the blinded review rankings, whereas runs from users with single-cell expertise (SC Experienced) exhibited variability and occasional lower-ranked outputs (Fig. 6b). Across all modes (HOTL, HITL, fully autonomous), outputs achieved a median credibility score of 2 (Fig. 6c; Supplementary Table 6; Methods), indicating that the outputs were deemed plausible but required stronger support. Interestingly, fully autonomous runs were perceived as having higher novelty scores (possibly new but need verification; Supplementary Table 6; Methods) while maintaining scientific credibility comparable to other interaction modes. In contrast, HITL interventions tended to recapitulate known biology with lower overall novelty scores, with most outputs receiving a median novelty score (Supplementary Table 6; Methods) below 2 (indicating no novelty). These results suggest that constant human intervention introduced a conservative bias to the reasoning process.

**Figure 6:**
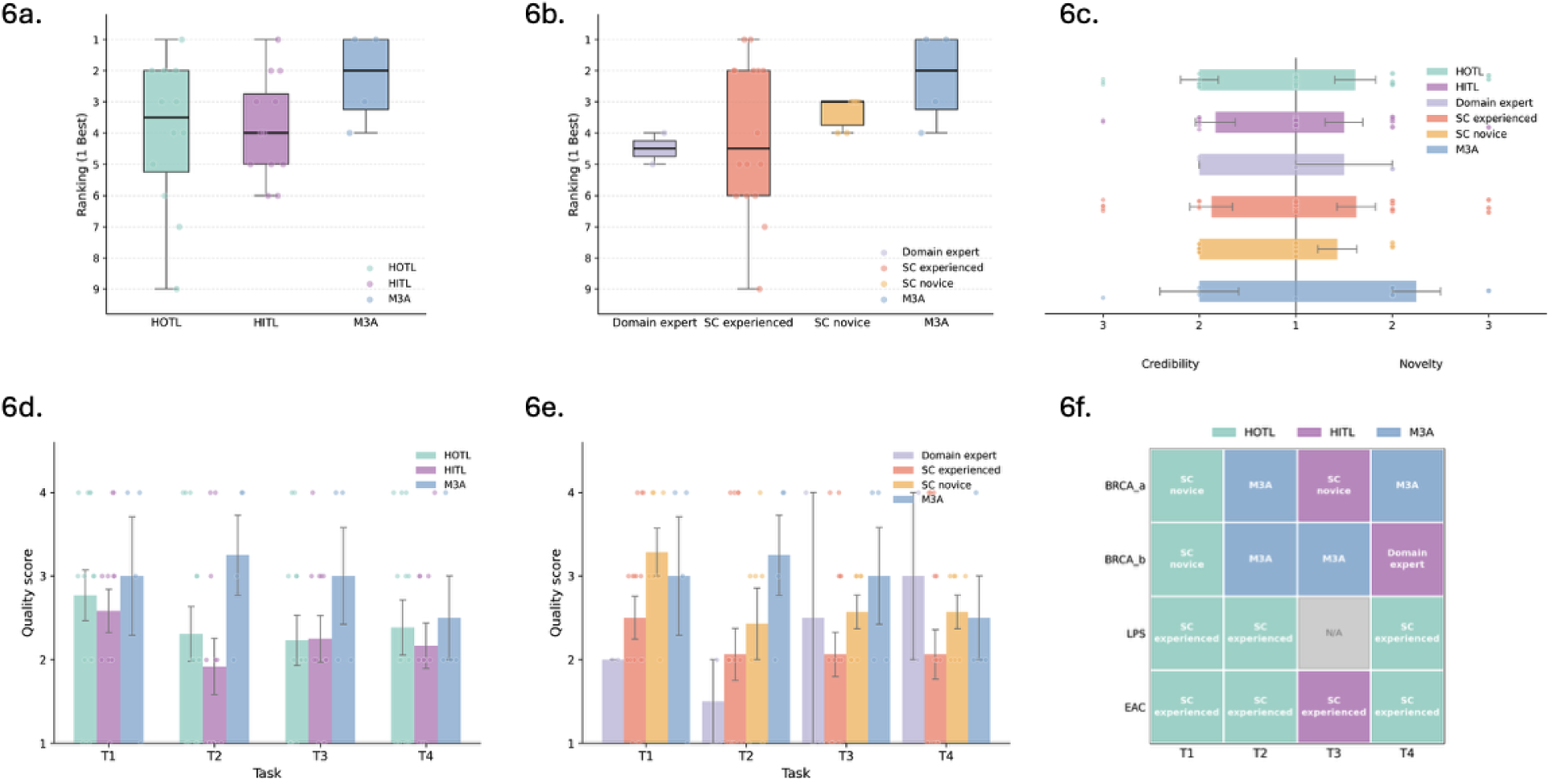
Blinded expert evaluation of human-AI copilot workflows across interaction modes and technical backgrounds. **(a-b)** Boxplots displaying the distribution of overall performance rankings (y-axis, where 1 is best) stratified by **(a)** interaction mode and **(b)** technical background, overlaid with individual data points representing independent rankings. Horizontal lines within boxes represent the median. **(c)** Diverging bar chart showing blinded expert ratings for Credibility (left) and Novelty (right) across both interaction modes and technical backgrounds. **(d)** Quality scores across four distinct biological reasoning tasks, stratified by human-AI interaction mode: Human-on-the-loop (HOTL, green), Human-in-the-loop (HITL, purple), and fully autonomous runs (blue). T1: cNMF program annotation, T2: Hypothesis formulation, T3: Regulatory driver analysis, T4: Biology synthesis. Error bars indicate standard deviation. **(e)** Quality scores for each task stratified by the user’s technical background: both single-cell and dataset-specific cancer biology expertise (Domain expert, light purple), single-cell expertise only (SC experienced, red), without prior single-cell experience (SC novice, yellow), and fully autonomous runs (blue). **(f)** Heatmap highlighting outputs that matched or were orthogonal to the original authors’ findings. Colored blocks indicate instances where the study authors ranked the output as equal or superior to their own for that specific task, labeled by the analyst’s background. Grey box indicates that no output was selected for the task.

Next, the blinded reviewers evaluated the quality of individual tasks with increasing biological complexity, spanning from foundational tasks (e.g. cNMF program annotation and hypothesis formulation) to more advanced tasks requiring sophisticated reasoning (e.g. regulatory driver analysis and biology synthesis tasks). Across all four biological reasoning tasks, most outputs were evaluated as adequate (median quality score of 2 [mostly correct; surface-level]) and fully autonomous modes consistently achieved quality scores comparable or superior to the human-AI copilot modes (HOTL and HITL) (Fig. 6d; Supplementary Table 6; Methods). Notably, fully autonomous mode evaluation scores progressively declined as task complexity increased. When stratifying performance by the human analyst’s background, workflows guided by individuals without single cell analysis backgrounds or autonomous execution yielded lower median quality scores in advanced reasoning tasks compared to early foundational tasks (Fig. 6e). While subjective biases initially influenced how blinded experts evaluated one another’s hypothesis formulations, this subjectivity progressively diminished in the more complex, open-ended biological tasks (regulatory driver analysis and biology synthesis), suggesting a potential benefit from the integration of single-cell and domain specific human expertise.

Lastly, we evaluated whether the copilot and autonomous workflows generated insights that matched or were orthogonal to the findings by the original study authors. Across the benchmark datasets, reviewers identified numerous tasks where workflows with minimal or no human involvement (HOTL or fully autonomous) produced outputs deemed superior or equivalent to their own original findings (Fig. 6f). Interestingly, for the more complex and reasoning-heavy regulatory driver analysis and biology synthesis tasks, reviewers favored the outputs generated with higher human intervention (HITL) or endorsed no output at all, underscoring that expert human steering remained important for complex biological reasoning.

## Discussion

Building upon growing enthusiasm for automated science and co-scientist efforts in the life sciences, evaluations of these AI agents have emphasized anecdotal use cases or tasks that largely focus on deterministic, workflow-automation^10,15–17^. Here, we investigated agentic life sciences research for open-ended discovery and biology reasoning with real-world cancer datasets, revealing both where automation is ready and where it is limited. Our results reveal current agentic AI is broadly competent at pattern recognition across well-charted biological territory, yet systematically constrained for discovery tasks.^20^

In the cell-type annotation tasks, we observed that AI annotation accuracy was associated with the density of training evidence rather than biological importance. For our multi-patient multi-cancer snMultiome datasets, concordance with expert labels was high for abundant cell populations, whose canonical marker genes are extensively documented, but failed disproportionately on rare cell types. We note that we did not provide a fixed cell-type vocabulary to the AI agents to select from, as annotation granularity varies across cancer types, datasets, and annotators. Instead, we developed a hierarchical grading schema to take into account these variations. This approach better reflects real-world workflows, where labels are not predefined and ontological guidance is often incomplete. While a constrained vocabulary would likely improve performance, it would make the evaluation less representative of atlas-scale disease annotation. Together, these findings suggest that benchmark performance on canonical cell types overestimates readiness for atlas-scale annotation in disease contexts, and that closing this gap will require targeted training on underrepresented biology rather than general model scaling.

At the level of biological reasoning, current AI-agents functioned in large part as knowledge-retrieval systems rather than hypothesis-generating reasoners^20^, and this distinction carries practical consequences for discovery. The agent’s systematic preference for proliferative, cell-cycle, and immune programs (themes that dominate biomedical literature) over lineage-specific and patient-specific signals may reflect the biases of the literature on which it was trained or other limitations related to the data modalities being explored. Because the programs most likely to yield mechanistic novelty are precisely those with the least prior documentation, AI-assisted prioritization may currently be most unreliable exactly where it could be most needed.

Our human-AI copilot experiment highlighted a potential model for future use of agentic AI that emphasizes the complementary strengths of AI agents and domain experts in the analysis of complex, real-world molecular datasets. Varying levels of human involvement and domain knowledge in our copilot experiments revealed a broad range of possibilities for AI-assisted biology discovery, with distinct effects on action allocation, technical depth, and the quality of biological interpretation. As biological reasoning tasks became more complex, our experiment also clarified the respective strengths and limitations of both agents and human experts.

While the M3A framework and the benchmarks described herein provide a foundation for systemic evaluation as agent capabilities evolve, our evaluation has several limitations. For the validation tasks utilizing the HTAN and TCGA datasets, one potential limitation is the inherent difficulty of cross-referencing cNMF programs derived from different data modalities (i.e., single-cell vs. bulk RNA-seq). Leveraging external single-cell datasets matched by specific cancer types may help reduce confounding variability. Regarding human-AI interaction assessments, the small cohort size and a single autonomous run per dataset required reporting of descriptive trends rather than statistical significance. Larger studies involving broader users may further inform the generalizability of the findings noted herein. Additionally, the concentration of human effort in early workflow stages may stem from temporal attention decay or novice users self-selecting out of advanced biological evaluations. The fixed experimental order, where users completed HOTL runs before HITL runs on the same dataset, means learning effects could have skewed subsequent behaviors. Our copilot experiment leveraged an expert-curated framework featuring pre-processed multi-omic datasets, biologically targeted prompts, and standardized output deliverables. Consequently, even fully autonomous runs inherently operated within a foundation of embedded human domain expertise. Despite these constraints, our multistep, multimodal, and multi-omic agentic evaluation framework provides important qualitative insights, highlighting that future frameworks must move beyond larger models to embrace architectures capable of explicit hypothesis revision over extended analytical horizons.

Ultimately, the M3A framework and our empirical findings point toward a paradigm shift in how AI is deployed in life sciences discovery. Rather than viewing autonomous agents and human oversight as competing modalities, the future of automated biological reasoning lies in a hybrid, sequential architecture: front-loading fully autonomous execution to exploit the agent’s capacity for scalable, systematic exploration, and transitioning to targeted HOTL or HITL interventions to anchor downstream biological interpretation in human judgment, contextual reasoning, and disciplinary expertise. By transforming AI agents from isolated execution engines into dynamic, context-aware collaborators, this hybrid approach lays the groundwork for a new era of accelerated scientific discovery, where the scale of machine computation is harmonized with the creative intuition of the human mind.

## Acknowledgements

S.J. was supported by the McDonough Fellowship in Metastatic Prostate Cancer Research and Artificial Intelligence. J.Y. was supported by the Eric and Wendy Schmidt Center at the Broad Institute of MIT and Harvard. J.P. was supported by NIH grant R50CA265182. E.V.A. was supported by grants P50CA272390, DOD HT94252410415.

## Author Information

These authors share senior authorship: Jihye Park, PhD, Eliezer Van Allen, MD.

## Corresponding authors

Correspondence to Eliezer Van Allen, MD

## Authors Contribution Statement

S.J., J.P., and E.V.A. conceived the study and designed the experiments. S.J. developed the code for the M3A framework. E.P., J.Y., S.C., and J.P. advised on the selection and inclusion of tools within the M3A framework. S.J. and J.P. performed data curation. E.P., J.Y., J.F., E.L.B., H.J., B.R., S.B., M.S., D.F., W.M., S.J., and J.P. participated in the copilot experiments. E.P., J.Y., and J.F. performed expert human annotations for the copilot experiments. S.J. and J.P. coordinated with the participants for the copilot experiments. S.J., J.P., and E.V.A. wrote the manuscript. All authors provided critical feedback on the manuscript, and approved its submission.

## Code and Data Availability

### Code availability

Code for M3A Agents is available at: https://github.com/vanallenlab/agentic-ai-codebase. Code for analysis is available at: https://github.com/vanallenlab/agentic-ai-analysis.

### Data availability

This study did not generate any new data. All data used in the study is publicly available. HTAN data used in the cell-type annotation and validation tasks can be downloaded from Terekhanova et al.^23^ (Pubmed ID: 37914932, DOI: 10.1038/s41586-023-06682-5). Data used in the copilot experiments can be downloaded from Pimenta et al.^27^ (Pubmed ID: 41564157, DOI: 10.1126/scitranslmed.adw4689) Yates et al.^28^ (Pubmed ID: 40499545, DOI: 10.1016/j.xcrm.2025.102188) and Fu et al.^29^ (Pubmed ID: 41315371, DOI: 10.1038/s41467-025-66659-y).

Data from HITL, HOTL and fully autonomous runs that was used for blinded expert grading can be found at https://drive.google.com/drive/folders/1V6rzANmdUg4miOtu5umKhpHC5rXW0LNM?usp=sharing.

## Methods

### Datasets

Four publicly available dataset collections were used for assessment of agentic-AI capabilities: Terekhanova et al.^23^ (HTAN-BRCA, HTAN-CESC, HTAN-CRC, HTAN-GBM, HTAN-HNSCC, HTAN-OV, HTAN-PDAC, HTAN-SKCM, HTAN-UCEC), Pimenta et al.^27^ (Pimenta-LPS), Yates et al.^28^ (Yates-EAC), and Fu et al.^29^ (Fu-BRCA). These datasets collectively spanned 11 different cancer types, over 182 patients and 1,064,365 cells. The published data from DFCI^27–29^ were analyzed as part of Institutional Review Board (IRB)-approved Dana-Farber Cancer Institute research protocol #26-243.

### Terekhanova et al.^23^ (HTAN-*)

HTAN is an atlas level dataset that was generated using consensus^30^. Patient and sample level information can be found in Supplementary Table 1. We retained solid tumor datasets with sufficient patient representation for downstream analysis. HTAN-MM was excluded as a non-solid tumor cohort, and HTAN-ccRCC was excluded because it included only a single patient and contributed too few cells. The final HTAN cohort comprised 646,057 sequenced cells from 146 samples across 130 patients, representing 9 cancer types.

### Pimenta et al.^27^

High-quality paired snMultiome dataset was generated from liposarcoma and normal adipose tissue obtained from 23 patients (Supplementary Table 1). Following quality control (QC) filtering, 22 snRNA (n=147,405 cells) and 19 snATAC (n=15,984 cells) samples were included for downstream analysis. Due to ATAC-specific QC thresholds, the final ATAC barcode was restricted to a strict subset of the verified RNA barcodes. Key clinical metadata was annotated for all samples, including sample id, pathology (WDLPS vs. DDLPS), treatment history (neoadjuvant_XRT), biological sex, age at resection, and anatomical biopsy site.

### Yates et al.^28^

High-quality paired snMultiome dataset was generated from primary and metastatic EAC samples obtained from 10 patients (n=21,444 cells; Supplementary Table 1). Following quality control (QC) filtering, one snATAC-seq library was excluded from downstream analysis (n=33,966 cells). Due to ATAC-specific QC thresholds, the final ATAC barcode was restricted to a strict subset of the verified RNA barcodes. Key clinical metadata was annotated for all samples, including sample id, tumor status (primary vs. metastatic), treatment history (therapy-naive vs. therapy-exposed), biological sex, and anatomical biopsy site.

### Fu et al.^29^

High-quality paired snMultiome dataset was generated from 40 breast cancer tumor biopsies obtained from 20 patients enrolled in a neoadjuvant immunotherapy and chemotherapy trial (snRNA: n=249,459 cells; snATAC: n=22,125 cells; Supplementary Table 1). This cohort comprised patients with HR+ (ER+/HER2-) breast cancer, with longitudinal sampling across multiple timepoints and two distinct treatment arms. 32 samples from 17 patients yielded paired snMultiome data, while 8 samples from 3 patients yielded snRNA-seq data only. Key clinical metadata was annotated for all samples, including sample id, PAM50, best response, timepoint, treatment_arm, treatment_timepoint, er_status, her2, and RCB index.

### TCGA^31^

The TCGA dataset was used as an held-out validation cohort for assessment of falsifiable. Patient and sample level information can be found in Supplementary Table 1.

### M3A framework

To rigorously evaluate how large language models (LLMs) navigate complex biological data, we developed M3A, an agentic evaluation framework. M3A is engineered to track and assess multi-step workflows, multimodal inputs, and multi-omics data integration (Fig. 1). Unlike static benchmarks, M3A maintains a controlled execution environment that records the model’s entire analytical trajectory, enabling both mechanistic insights and quantitative benchmarking. Consistent and interpretable evaluation of agentic scientific reasoning requires control over five core dimensions: the execution environment, the tool suite, multimodal context integration, persistent data state, and step-level telemetry.

### Standardized execution environment

All agent executions run within a standardized computational environment initialized from a common Docker image. Agents may install dependencies and modify the environment during a run as needed, but all such changes are reset between runs. This design ensures consistent initial conditions across evaluations while preserving the flexibility required for complex analyses, eliminating cross-run contamination and supporting reproducible benchmarking.

### Specialized tool suite

M3A equips agents with four categories of tools. Python-based programmatic analysis supports data exploration and manipulation, development of novel analytical methods, and flexible non-standard processing driven by the agent’s own reasoning about the data. Terminal access allows agents to install packages not present in the default environment, though it is otherwise restricted to preserve execution safety. Domain-specific tools for snRNA-seq and snATAC-seq provide curated, easy-to-use functions covering individual steps of standard single-cell processing workflows. Finally, web-based literature retrieval enables agents to query Google Search and domain-specific repositories, including PubMed, arxiv and Google Scholar, to ground analytical decisions in published research. Anthropic’s native search tool is also included. To prevent data leakage during experimental runs, both the web-search tool and Anthropic’s native search tool were configured to blacklist URLs associated with the published study, including the source manuscript, code repository, and manuscript assets such as data availability links.

### Multimodal context integration

M3A supports long-horizon reasoning by recursively incorporating model-generated outputs, including natural language, code, and visualizations, into an evolving context window at each step. This recursive accumulation of multimodal outputs allows agents to iteratively build on prior results, maintaining coherence across extended analytical workflows.

### Persistent data state

Structured data objects, such as AnnData instances, either declared during initialization of the agent or generated during the reasoning and analysis, persist across tool invocations throughout a session. This persistent state preserves intermediate computational results, enabling agents to reference and extend prior steps of the analyses, a prerequisite for coherent multi-step biological analyses.

### Step-level telemetry

M3A captures the full sequence of agent decisions through comprehensive step-level instrumentation. Before each action, the system records the model’s stated intent, tool selection rationale, and anticipated outcomes via pre-tool call hooks. Execution traces log generated outputs, visualizations, token usage, and structured interaction data are logged automatically on wandb. Optional think-aloud voice notes in case of co-pilot experiments are captured via AI-generated meeting summaries using zoom’s inbuilt tool to provide additional visibility into intermediate reasoning. In addition, users recorded optional, real-time notes throughout the copilot sessions to provide qualitative context. This instrumentation enables direct analysis of decision trajectories, including how errors propagate and how agents recover, across complex workflows.

### Autonomous AI-driven cell-type annotation

#### Cohorts and reference labels

We compared AI-generated cell-type annotations against expert-curated ground truth (GT) from the Terekhanova et al.^23^ across nine tumor cohorts (BRCA, CESC, CRC, GBM, HNSCC, OV, PDAC, SKCM, UCEC). For each cell, the AI agent generated a single free-text label. The author annotations contained fine-grained labels (GT_fine). Broad-category labels (GT_broad) were derived automatically from each cohort’s cell-type ontology by traversing the parent chain from any GT_fine node to the root, yielding six top-level lineages: epithelial, stromal, endothelial, immune, neural, and cycling. Cells whose GT label was flagged as low quality, doublet, or unknown were excluded from scoring, as were cells whose GT label lacked an entry in the cohort ontology; both categories are combined in the denominator when reporting schema coverage.

#### Label-mapping schemas

For each cohort, a hierarchical cell-type ontology was constructed as a rooted tree in which every node represents a cell-type concept at a defined resolution. Each observed AI label was mapped to a single ontology node, as was each GT_fine label. Ontologies were required to cover every AI label observed in the cohort; any unmapped AI label was treated as an error. The tree structure encodes biological resolution: a node’s ancestors represent coarser cell-type concepts and its descendants represent finer ones. Classification of each cell was performed by locating the AI node and the GT node in the tree and applying the 6-condition scheme described below.

#### Tier classification

Each scored cell was assigned to one of six tiers.

- Exact match (c1): the AI node and GT node are identical.
- Lineage-correct (c2): the AI node is a strict ancestor of the GT node, and no other AI vocabulary term maps to the GT node or any of its ontology descendants (i.e., the AI could not have been more specific given its labeling vocabulary).
- More specific than GT (c6): the AI node is a strict descendant of the GT node.
- Sibling miss (c3): the AI node and GT node share the same immediate parent but are not identical.
- Specifiable miss (c5): the AI node is a strict ancestor of the GT node, but at least one other AI vocabulary term maps to the GT node or a descendant of it, meaning a finer label was available and not used.
- Lineage error (c4): none of the above apply; the AI and GT nodes fall in different branches of the ontology.

#### Cohort-level metrics

The accuracy was computed as (n_c_1_ + n_c_2_ + n_c_6_) / n_scored, reflecting credit for exact, lineage-correct, and more-specific-than-GT placements. Fine accuracy was defined as (n_c_1_ + n_c_6_) / n_scored, capturing only unambiguous subtype-level matches. Schema coverage was defined as n_scored / n_total, where n_total includes all cells in the AI-GT intersection and n_scored excludes those with excluded or unmapped GT labels.

#### Confidence intervals

Per-cohort 95% confidence intervals were obtained by nonparametric bootstrap (n=1,000 resamples), resampling HTAN sample identifiers with replacement to preserve within-sample correlation. GBM, which contained only two samples, was bootstrapped at the cell level.

#### Macro sensitivity and specificity

Cohort-level sensitivity and specificity were computed at the broad lineage tier using a one-vs-rest decomposition of the multi-class confusion matrix, treating each broad lineage class as equally important regardless of cell count. Predicted broad labels were obtained by mapping each AI ontology node to its top-level lineage using the same tree-walk used for GT_broad. For each class present in the cohort, standard true-positive, false-negative, false-positive, and true-negative counts were derived from the predicted versus ground-truth broad-label assignments. Per-class sensitivity and specificity were then averaged uniformly across classes to yield cohort-level macro scores. Because true negatives dominate per-class denominators when any individual lineage comprises a small fraction of total cells, specificity is structurally high across all cohorts and is reported primarily as a sanity check; sensitivity is the operationally informative summary statistic.

### Prioritization Tasks

#### Design Principles

Validation tasks were designed to test whether transcriptional programs discovered from single-cell data recapitulate known clinical and molecular phenotypes in independent bulk cohorts. Tasks were defined by pairing a cancer type with a clinically or biologically meaningful endpoint derived from the TCGA data sources described above. Three selection criteria governed task inclusion:

#### Prevalence

Each candidate endpoint was screened for sufficient prevalence to support statistical testing. For binary endpoints (e.g., mutation status, receptor positivity), a minimum minority-class frequency was required (typically >10%). For survival endpoints, a minimum event rate of 20% was required. For continuous and ordinal endpoints, a minimum of approximately 50 labeled samples was required. Endpoints failing these thresholds were excluded or flagged as exploratory.

#### Endpoint independence from expression

Endpoints were chosen to be orthogonal to the RNA-seq data used for program activity scoring. Validation labels were derived from DNA-level alterations (somatic mutations, copy number, methylation), protein measurements (RPPA, IHC), histopathologic features (LLM-extracted from pathology reports), and censored survival. None of these data values were used in cNMF program discovery or ssGSEA scoring.

#### Clinical relevance

Tasks were anchored to established clinical or molecular phenotypes with therapeutic, prognostic, or subtyping significance for each cancer type (e.g., receptor status in breast cancer, MSI/MMR deficiency in colorectal cancer, HRD in ovarian cancer).

Tasks spanned three categories (molecular, pathology, and clinical), and included both cancer-type-specific endpoints and pan-cancer endpoints applied across eligible cohorts (see Supplementary Table 3). Example prompts can be found in Supplementary Note 3.

### Endpoint extraction

#### Molecular data

Molecular data for 11 TCGA cohorts were obtained from cBioPortal ^32,33^ (GDC 2025 release, reference genome GRCh38/hg38; source data generated via the Cancer Data Aggregator from the NCI Genomic Data Commons). The cohorts comprised breast invasive carcinoma (BRCA; n=1,102), clear cell renal cell carcinoma (ccRCC; n=537), cervical squamous cell carcinoma and endocervical adenocarcinoma (CESC; n=309), colon adenocarcinoma (COAD; n=463), rectal adenocarcinoma (READ; n=171), glioblastoma multiforme (GBM; n=611), high-grade serous ovarian carcinoma (HGSOC; n=604), head and neck squamous cell carcinoma (HNSC; n=530), pancreatic adenocarcinoma (PAAD; n=186), skin cutaneous melanoma (SKCM; n=474), and uterine corpus endometrial carcinoma (UCEC; n=549). For each cohort, RNA-seq gene expression (TPM, FPKM, and read counts), somatic mutations (MAF format), discrete copy number alterations (GISTIC 2.0), segmented copy number data were obtained. Additionally, DNA methylation beta values and reverse-phase protein array (RPPA) data were obtained from TCGA PanCancer Publications ^34^. COAD and READ were analyzed jointly as COADREAD for colorectal-specific tasks.

#### Somatic mutations (DNA)

Tumor and matched normal samples were profiled by whole exome sequencing (WES). Reads were aligned to GRCh38 and somatic variants were called by the GDC DNA-Seq pipeline using an ensemble of four callers: MuTect2, MuSE, VarScan2, and Pindel. Variants were reported in Mutation Annotation Format (MAF) with variant classification, reference/alternate allele counts, and protein-level effect annotations (HGVSp_Short).^35,36^

#### Copy number alterations (DNA)

Tumor and matched normal DNA were profiled on the Affymetrix Genome-Wide Human SNP Array 6.0. Raw signal intensities were segmented by circular binary segmentation (CBS) using the DNAcopy R package, producing genomic segments with log2 copy number ratios (tumor minus matched normal). Gene-level discrete calls were generated by the GISTIC 2.0 algorithm: −2 (homozygous deletion), −1 (hemizygous deletion), 0 (neutral), 1 (gain), 2 (high-level amplification). Both segmented and discrete data were used in downstream analyses. ^37,38^

### DNA Methylation

Genomic DNA was bisulfite-converted (unmethylated cytosines converted to uracil; methylated cytosines unchanged), amplified, and hybridized to the Illumina Infinium HumanMethylation450 (450K) BeadChip array, which interrogates >480,000 CpG sites covering 96% of CpG islands. Methylation levels were quantified as beta values, computed as M/(M+U+100), where M and U are methylated and unmethylated signal intensities and 100 is a regularization constant. Pan-cancer data were obtained from the JHU-USC 450K dataset. ^39^

### Bulk RNA-seq

Total RNA was sequenced on Illumina HiSeq platforms. Reads were aligned to GRCh38 using the STAR two-pass method (GDC mRNA analysis pipeline) and gene-level expression was quantified by STAR counts using GENCODE gene annotations. Expression values were reported as transcripts per million (TPM), fragments per kilobase of transcript per million mapped reads (FPKM), and raw read counts, indexed by Entrez Gene ID. ^39–41^

### Reverse-phase protein assay (RPPA)

Protein lysates from tumor samples were printed onto nitrocellulose-coated slides and probed with 198 validated antibodies targeting total, phosphorylated, and cleaved protein forms. Protein expression was estimated by SuperCurve fitting from 5-point serial dilution series, and replicate-based normalization (RBN) was applied across batches. Values were reported as median-centered, normalized log2 protein expression levels. ^42–44^

### Clinical and survival data

Time-to-event data were obtained from the TCGA Clinical Data Resource (TCGA-CDR), which provides uniformly curated overall survival (OS), disease-specific survival (DSS), disease-free interval (DFI), and progression-free interval (PFI) with event indicators and follow-up times in days for 11,160 patients across 33 TCGA cancer types. Patients were mapped to cancer cohorts via the CDR ‘typè field, and records with missing event status or non-positive follow-up times were excluded. Survival endpoints were selected per cancer type based on event rate viability (minimum 20% event rate) ^31^.

Clinical biomarker annotations were obtained from the TCGA pan-cancer clinical follow-up file, which aggregates site-reported clinical data across all TCGA studies. Fields used included estrogen receptor (ER) status, progesterone receptor (PR) status, HER2 IHC and FISH status (BRCA only), microsatellite instability (MSI) status, and HPV status by p16 testing. Records with sentinel missing values (Not Available, Not Evaluated, Unknown, Discrepancy) were excluded.

### Pathology reports

A total of 9,523 TCGA surgical pathology reports were obtained from the TCGA-Reports. The original TCGA pathology reports, present as PDFs, were processed to remove formatting artifacts, producing a corpus of patient barcode--report text pairs. ^45,46^

### Report-to-cohort assignment

Reports were matched to the 11 TCGA cohorts by extracting the 12-character patient barcode (TCGA-XX-XXXX) from each report filename and cross-referencing against the case lists for each cBioPortal cohort. This assigned 4,530 reports to a cancer type: BRCA (n=1,031), ccRCC (n=524), CESC (n=289), COAD (n=418), READ (n=161), GBM (n=397), HGSOC (n=366), HNSC (n=520), PAAD (n=176), SKCM (n=102), and UCEC (n=546). The remaining 4,993 reports did not match any cohort and were excluded.

### Schema-guided structured extraction

For each cancer type, a structured JSON extraction schema was developed defining every histopathologic feature to extract from the free-text reports. Schema construction followed a two-phase protocol. In Phase A (vocabulary research), three rounds of iterative corpus sampling (30 reports per round, non-overlapping, with fixed random seeds) were conducted per cancer type, preceded by a corpus-wide n-gram frequency analysis. Each round cataloged synoptic field:value patterns, prose phrasings, clinical abbreviations, OCR artifacts, international naming variants, and “not specified” phrasings; the vocabulary was iteratively expanded until saturation. In Phase B (schema output), the frozen vocabulary was encoded into a JSON schema. Each field was typed as binary, categorical, or continuous, with indicator lists for each value state (minimum 10 entries per indicator list for binary fields). Universal fields common to all cancer types included histologic type, histologic grade, tumor size, surgical margin status, lymphovascular invasion, perineural invasion, lymph node status, necrosis, mitotic rate, and pathologic TNM staging. Cancer-specific fields were added per cancer type (e.g., Nottingham sub-scores for BRCA, Fuhrman grade for ccRCC, POLE ultramutator markers for UCEC).

### LLM-based extraction

Structured field extraction was performed using Claude Sonnet-4.6^47^. Each report was sent as a user message (see Supplementary Note 3) with a system prompt encoding the full schema, field definitions, indicator lists, and missing data policy. The model returned a structured JSON object with one value per schema field. The missing data policy distinguished four states: present/positive, absent/negative, null (not mentioned), and “not_specified” (acknowledged but not assessed).

### Program activity scoring and statistical validation

#### Transfer of cNMF programs to bulk RNA-seq

Gene programs discovered by consensus non-negative matrix factorization (cNMF) on HTAN scMultiome data were used to score TCGA bulk RNA-seq using single-sample gene set enrichment analysis (ssGSEA)^48^. For each cNMF program, the top contributing genes were mapped to Entrez gene IDs via the Hugo-to-Entrez mapping derived from cBioPortal expression matrices. Per-sample ssGSEA enrichment scores were then computed from the TPM expression matrix, producing a programs-by-samples activity matrix for each cohort.

#### Validation metric and ranking

Each validation task tested the association between every program’s ssGSEA activity score and the task’s clinical/molecular endpoint. The validation metric was chosen based on endpoint type: AUROC for binary endpoints for axis-free tasks such as histotype classification), Spearman rank correlation for continuous and ordinal endpoints, and the absolute z-score from a univariate Cox proportional hazards model for time-to-event endpoints. Programs were ranked by their metric value within each task, producing a ground-truth statistical ranking. The top-ranked program represents the strongest statistical association between a discovered transcriptional program and the clinical endpoint.

#### Concordance analysis: endpoint-derived vs. agent-generated rankings

For each validation task, two independent rankings of the same set of cNMF programs were compared: the endpoint-derived ranking, produced by a statistical validation procedure (programs ordered by measured association strength with the clinical/molecular endpoint), and the agent-generated ranking, produced by an M3A agent that ranked programs based on biological interpretation of the gene lists without access to the TCGA endpoint data. For each task, the two rankings were aligned on the common set of programs. Each program received a pair of ranks: its position in the endpoint-derived ranking and its position in the agent-generated ranking. Agreement between the two rankings was quantified at multiple levels:

- *Top-1 accuracy:* whether both rankings identified the same program as rank 1.
- *Top-3 swap:* the number of programs appearing in both rankings’ top 3 (out of 3).
- *Spearman rank correlation:* the full-ranking correlation between endpoint-derived and agent-generated ranks across all programs in a task.

To contextualize the observed concordance, permutation baselines were computed for each task. For each distinct number of programs (N), 10,000 random permutations were generated and the same concordance metrics (top-1 accuracy, top-3 overlap) were computed. Baselines were then weighted by the number of tasks at each N to produce an overall expected-by-chance reference.

Concordance metrics were aggregated and compared across task category (molecular, pathology, clinical), cancer type, validation metric type (AUROC, Spearman, Cox), and endpoint signal strength (within-metric-type percentile rank of the best program’s score).

### Copilot Experiments

#### Approach

The human-AI copilot was conducted in two formats amongst 11 participants, and all participants completed both, except one participant only completed HOTL. In the human-on-the-loop (HOTL) format, the agent ran autonomously while the human monitored progress; however, users were instructed to intervene only upon encountering major analytical disruptions, such as model hallucinations or clearly erroneous reasoning. In the human-in-the-loop (HITL) format, the human was an active participant who could request additional analyses, plots, or other actions at any point during execution. Participants completed the HOTL run first, as this provided familiarity with the agent’s autonomous behavior and informed decisions about when and how to intervene during the HITL run.

At each step of the HITL run, the agent paused and awaited a human response before proceeding. Participants selected from four response options. “Continue” instructed the agent to proceed without intervention. “Feedback” allowed participants to provide direct instructions — including requests for plots, repetition of an analysis step, database queries, step-level explanations, general guidance for subsequent steps, or simply directing the agents to proceed to the next phase of the workflow; all feedback was retained in the agent’s context. “Questions” allowed participants to pose clarifying questions without the agent retaining the question or response in memory; this option was used sparingly. “Abort” terminated the run and was reserved for cases in which the agent had entered an unrecoverable loop or exhibited significant, sustained hallucinations across multiple steps.

To capture holistic thoughts of the participants, part of which may not be conveyed through written instructions to the agent, participants joined a Zoom meeting during the experiment to capture AI-generated meeting summaries or recorded real-time notes throughout the copilot sessions.

#### Experiment Design

We defined three distinct human-AI interaction modes in this framework: 1) Human-in-the-loop (HITL), where humans actively steer the AI agent’s reasoning by providing real-time feedback and decision-making; 2) Human-on-the-loop (HOTL), where humans monitor the AI agent’s actions and intervene only to correct errors or enforce constraints, without directing its reasoning process; and 3) Autonomous (M3A), where the AI agent operates independently without any human oversight.

To evaluate the framework across diverse biological contexts, we utilized the three published datasets^27,28,29^, each associated with the specific biological questions addressed in their original studies. LPS^27^ datasets were used to characterize the transcriptomic and epigenetic features that are different between well-differentiated (WDLPS) and dedifferentiated (DDLPS) liposarcoma subtypes. EAC^28^ datasets are focused to identify malignant cell programs, both transcriptomic and epigenetic, associated with the progression of esophageal adenocarcinoma. Lastly, BRCA^29^ datasets were utilized to determine the malignant cell programs that drive therapy resistance in HR+ breast cancer patients treated with chemo-immunotherapy.

In order to evaluate how varying levels of expertise influence AI agent utility within standard biological workflows, we asked a representative cohort of computational biologists to analyze each dataset. Participants were categorized into four groups based on their technical and domain-specific experience: 1) The first authors of the studies or dataset experts with both single-cell expertise and dataset-specific cancer biology domain knowledge (Domain expert); 2) Researchers with one or more years of active single-cell RNA-seq project experience (SC experienced); 3) Researchers with no single-cell analysis experience or fewer than six months of exposure (SC novice); and 4) Autonomous AI agent (M3A), serving as a baseline for non-human intervention.

Each participant performed four biological reasoning tasks of increasing complexity, reflecting the standard analytical and interpretative workflows of a computational biologist. Task 1 (cNMF program annotation) tasked to assign biological labels to cNMF programs based on top weighted features with a structured justification; Task 2 (Hypothesis formulation) required the agent to rank the two candidate programs most relevant to dataset-specific biological question and to identify patterns that would falsify the hypothesis. Task 3 (Regulatory driver analysis) focused to identify candidate regulatory drivers for the top-ranked program from Task 2, requiring both a methodological approach and a list of specific candidate genes. Finally, Task 4 (Biology synthesis) required the agent to synthesize previous tasks into a discovery model, connecting the regulatory architecture (Task 3) to the transcriptional programs (Task 1) to support the biological hypothesis (Task 2). Scientists were asked to submit their log files with all their interactions with the agent and final outputs, including a summarized write-up per task from each interaction mode (HOTL and HITL) (Data availability).

To ensure a rigorous and standardized evaluation, we explicitly pre-configured our copilot environment with extensive domain expertise. First, input datasets were comprehensively pre-processed to integrate multi-omic profiles, specifically scRNA and scATAC data, alongside cNMF usage scores, broad cell-type annotations, and relevant clinical metadata. Second, prompts were engineered to encode dataset-specific objectives and targeted biological questions for each dataset (Supplementary Note 3). Finally, output deliverables were strictly standardized to enable uniform, quantitative downstream evaluation. We ensured that human domain expertise served as a foundational constraint within the experimental design, systematically guiding even the fully autonomous runs.

#### Evaluation Design

The workflows and final outputs were subsequently evaluated using two strategies: 1) human and AI’s actions were classified into distinct functional categories to track technical breadth and guidance; and 2) the first authors of the originating studies conducted a blinded expert review of the unlabeled deliverables.

To characterize the technical breadth of each session, we classified agent actions into nine functional categories:

- Data exploration: Actions dedicated to inspecting data structures, dimensions, column names, value distributions, metadata, and object attributes. (e.g., executing head(), tail(), or checking for missing values).
- Data wrangling: Operations involving data cleaning, subsetting, filtering, merging, reshaping, or normalization, alongside feature engineering and missing-value imputation. This category also encompassed environment configuration, including package installations (pip install), library imports, and directory management.
- Domain-specific bioinformatics: Execution of computational pipelines requiring biology-specific parameters or data structures (e.g., AnnData objects, peak-by-cell matrices). This included differential expression analysis (DESeq2/Wilcoxon), gene set enrichment (GSEA/gseapy), cNMF, trajectory inference, cell-type annotation, chromVAR, copy-number inference, and peak calling.
- Result interpretation: Isolated cognitive assessments where the agent explained or labeled a single finding (e.g., assigning a cell-type identity based on marker genes) utilizing its pre-trained knowledge base without external tool queries.
- Literature search: Dynamic queries executed against external databases or web resources (e.g., PubMed, reference repositories) to retrieve biological annotations or external scientific context.
- Statistical testing: Domain-agnostic statistical procedures applied without requiring specific biological context, including t-tests, Mann-Whitney U tests, chi-squared tests, ANOVA, regression modeling, and principal component analysis (PCA).
- Visualization: The generation, formatting, and saving of any graphical figures or plots, such as scatter plots, UMAPs, heatmaps, volcano plots, and violin plots.
- Cross-analysis synthesis: Complex analytical steps requiring the explicit integration of findings from two or more prior, distinct subanalyses (e.g., synthesizing cNMF program annotations with ATAC motif enrichments) into a unified narrative or overarching biological hypothesis.
- Error recovery: Targeted debugging actions where the primary objective was diagnosing and resolving an explicit exception or unexpected tool failure from a preceding step, including traceback inspection and parameter adjustments.

We also classified all user interventions into ten functional categories:

- Method direction: Explicit instructions dictating the methodological approach, analytical technique, or specific software tool the agent should deploy for a given task (e.g., specifying a particular enrichment package or data subset).
- Requesting deliverable: Directives requiring the agent to synthesize analytical outputs into a formalized work product, such as a written summary, hypothesis formulation, or structured biological report, distinct from simple progress check.
- Refocusing: Corrective interventions deployed when the agent deviated from the desired analytical path, performed redundant actions, or executed erroneous code, explicitly forcing the agent to abort or modify its current trajectory.
- Progress check: Diagnostic queries used to monitor agent status, verify task completion, or prompt the agent to justify its immediate reasoning without providing active guidance or direction.
- Task navigation: Neutral workflow coordination managing the operational sequence, including routine affirmations (e.g., “OK,” “proceed”) to signal progression to the next milestone when the agent is performing correctly.
- File management: Instructions regarding output directory mapping, path corrections, or file storage requirements, typically used to correct or specify where analytical assets should be saved.
- Control output verbosity: Directives manipulating the formatting, length, or granularity of the agent’s textual or computational responses (e.g., requesting concise summaries instead of raw code dumps).
- Domain knowledge: Proactive provision of specialized scientific context, biological facts, experimental design nuances, or dataset metadata intended to anchor or correct the agent’s analytical grounding.
- Constraint setting: The imposition of standing behavioral rules or global limitations designed to restrict the agent’s operational parameters across multiple subsequent steps (e.g., forbidding package installations or enforcing global sample filters).
- Persona calibration: Informational prompts defining the user’s background, technical expertise, or pedagogical preferences to dynamically adjust the agent’s communication style and level of explanation.

Finally, we conducted a blinded expert review performed by the original study authors (Supplementary Table 6). BRCA^29^ datasets were reviewed by two domain experts. To ensure an unbiased assessment, reviewers were presented with the final outputs from copilot (HITL and HOTL) and fully autonomous runs without labels indicating the interaction mode or their technical background. In addition, we provided a standardized summary of analytical steps to contextualize the process without revealing human intervention (Data availability). Reviewers evaluated the quality of the four biological reasoning tasks based on scientific accuracy, depth of insight, and use of evidence. Beyond individual tasks, reviewers also provided the credibility and novelty scores, as well as a comparative ranking of all outputs. To measure the AI’s real-world utility, reviewers identified any outputs they considered equal or superior to their original findings. Finally, reviewers attempted to infer the interaction mode and the analyst’s level of expertise for each run.

## Supplementary Information

### Supplementary Notes

**Supplementary Note 1:** Cell-Type Ontologies (per cohort)

**Supplementary Note 2:** Complete log from a M3A agent run for cell-type annotation in melanoma (SKCM)

**Supplementary Note 3:** M3A agent’s system prompts used across different experiments (cell-type annotation, held-out validation tasks and copilot experiments).

### Supplementary Tables

**Supplementary Table 1**: Summary of Datasets.

**Supplementary Table 2**: Pan-cancer evaluation of AI-agent’s cell-type annotations against author annotations across nine cancer-types

**Supplementary Table 3**: Concordance between M3A agent program rankings and statistical ground-truth rankings

**Supplementary Table 4**: User behavior in copilot experiments

**Supplementary Table 5**: Effect of interaction mode and user experience on steering of M3A agents

**Supplementary Table 6**: Blinded expert review survey questions

### Supplementary Figures

**Supplementary Figure 1**: Per-cluster scrutiny depth versus broad annotation accuracy, stratified by cancer type (BRCA, CESC, CRC, GBM, HNSCC, OV, PDAC, SKCM, UCEC) and cell-type lineage (epithelial, stromal, endothelial, immune, neural, cycling).

**Supplementary Figure 2**: M3A agent scrutiny depth, annotation accuracy, and workflow strategy (scATAC usage, label transfer, gene activity scoring, UMAP concordance, quantitative metrics, copy-number variation (CNV) analysis, knowledge driven marker analysis, iterative re-analysis) by cancer type (OV, SKCM, PDAC, UCEC, HNSCC, GBM, CRC, CESC, BRCA).

**Supplementary Figure 3: (a)** Representative tasks from each quadrant of the signal-strength × M3A-agent’s outcome space (M3A hit, M3A miss); light-gray bar indicates the total number of cNMF programs in that task. **(b)** Representative example tasks of M3A-agent prioritization being an exact match (green, rank 1), near miss (orange, rank 2-3), or clear miss (red, rank ≥ 4) of the statistically top-1 program; light-gray bar indicates the total number of cNMF programs in that task.

**Supplementary Figure 4: (a)** M3A agent action category (data exploration, data wrangling, visualization, statistical testing, domain-specific bioinformatics, error recovery, result interpretation, literature search, cross-analysis synthesis) composition between AI-only and human+AI modes, stratified by tasks (Data exploration; Program annotation, T1; Hypothesis formulation, T2; Regulatory driver analysis, T3; Biology synthesis, T4). **(b)** Co-occurrence of alternative M3A agent actions.

**Supplementary Figure 5**: Impact of interaction mode (HOTL - HITL) in user interaction category (task navigation, output formatting, method direction, constraint setting, correcting/refocusing, requesting deliverable, file/output management, progress check, domain knowledge, personal calibration) composition, stratified by task stage (Program annotation, T1; Hypothesis formulation, T2; Regulatory driver analysis, T3; Biology synthesis, T4) and experience level (Domain Expert (DE), SC Experienced(SCE), SC Novice(SCN)).

**Supplementary Figure 6**: Reviewer identification of **(a)** interaction mode and **(b)** analyst background. Each bar represents a true interaction style (HOTL or HITL) or a true analyst background category (Domain expert, SC experienced, SC novice, M3A) and shows the distribution of reviewer guesses across all cancer cohorts pooled.

## References

1. Ghareeb, A. E. et al. A multi-agent system for automating scientific discovery. Nature 1–3 (2026).

2. Aygün, E. et al. An AI system to help scientists write expert-level empirical software. Nature 1–3 (2026).

3. Gottweis, J. et al. Accelerating scientific discovery with Co-Scientist. Nature 1–3 (2026).

4. Lu, C. et al. Towards end-to-end automation of AI research. Nature 651, 914–919 (2026).

5. Gao, S., et al. Democratizing AI scientists using ToolUniverse. (2025).

6. Huang, K., et al. Biomni: A General-Purpose Biomedical AI Agent. *bioRxiv* (2025) doi:10.1101/2025.05.30.656746.

7. Ouyang, L. et al. Training language models to follow instructions with human feedback. (2022).

8. DeepSeek-AI et al. DeepSeek-R1: Incentivizing Reasoning Capability in LLMs via Reinforcement Learning. (2025) doi:10.1038/s41586-025-09422-z.

9. Hubert, T. et al. Olympiad-level formal mathematical reasoning with reinforcement learning. Nature 651, 607–613 (2025).

10. Laurent, J. M., et al. LAB-Bench: Measuring Capabilities of Language Models for Biology Research. (2024).

11. Jin, Q., Yang, Y., Chen, Q. & Lu, Z. GeneGPT: augmenting large language models with domain tools for improved access to biomedical information. Bioinformatics 40, (2024).

12. Liu, H., Chen, S., Zhang, Y. & Wang, H. GenoTEX: An LLM Agent Benchmark for Automated Gene Expression Data Analysis. (2024).

13. Fa, D., Culjak, M., Pandza, B. & Cupic, M. BioAgent Bench: An AI Agent Evaluation Suite for Bioinformatics. (2026).

14. Website. doi:10.1101/2025.01.23.634608.

15. Alber, S. et al. CellVoyager: AI CompBio agent generates new insights by autonomously analyzing biological data. Nat Methods 23, 749–759 (2026).

16. Mao, Y., et al. scAgent: Universal Single-Cell Annotation via a LLM Agent. (2025).

17. Workman, K., Yang, Z., Muralidharan, H., Abdulali, A. & Le, H. scBench: Evaluating AI Agents on Single-Cell RNA-seq Analysis. (2026).

18. Website. https://www.biorxiv.org/content/10.64898/2026.04.29.721531v1.full.

19. Website. https://www.biorxiv.org/content/10.1101/2025.04.03.646459v1.

20. Mitchener, L. et al. Kosmos: An AI Scientist for Autonomous Discovery. (2025).

21. Website. doi:10.64898/2026.04.06.716850.

22. [No title]. https://www-cdn.anthropic.com/6a5fa276ac68b9aeb0c8b6af5fa36326e0e166dd.pdf.

23. Terekhanova, N. V. et al. Epigenetic regulation during cancer transitions across 11 tumour types. Nature 623, 432–441 (2023).

24. Rozenblatt-Rosen, O. et al. The Human Tumor Atlas Network: Charting Tumor Transitions across Space and Time at Single-Cell Resolution. Cell 181, 236–249 (2020).

25. de Bruijn, I. et al. Sharing data from the Human Tumor Atlas Network through standards, infrastructure and community engagement. Nature Methods 22, 664–671 (2025).

26. Kotliar, D. et al. Identifying gene expression programs of cell-type identity and cellular activity with single-cell RNA-Seq. (2019) doi:10.7554/eLife.43803.

27. Pimenta, E. M. et al. Epigenetic dysregulation of metabolic programs mediates liposarcoma cell plasticity. Science Translational Medicine (2026) doi:10.1126/scitranslmed.adw4689.

28. Yates, J. et al. Cell states and neighborhoods in distinct clinical stages of primary and metastatic esophageal adenocarcinoma. Cell Rep Med 6, 102188 (2025).

29. Fu, J. et al. Cellular reprogramming during anti-PD-1 and chemotherapy treatment in early-stage primary hormone receptor-positive breast cancer. Nature Communications 16, 10704 (2025).

30. HTAN Phase 2 Data Model — HTAN Phase 2 Data Model 0.1.0 documentation. https://htan2-data-model.readthedocs.io/en/main/index.html.

31. Liu, J. et al. An Integrated TCGA Pan-Cancer Clinical Data Resource to Drive High-Quality Survival Outcome Analytics. Cell 173, 400–416.e11 (2018).

32. Cerami, E. et al. The cBio cancer genomics portal: an open platform for exploring multidimensional cancer genomics data. Cancer Discov 2, 401–404 (2012).

33. Gao, J. et al. Integrative analysis of complex cancer genomics and clinical profiles using the cBioPortal. Sci Signal 6, l1 (2013).

34. TCGA - PanCanAtlas Publications. https://gdc.cancer.gov/about-data/publications/pancanatlas.

35. Bioinformatics Pipeline: DNA-Seq Analysis: Whole Exome and Targeted Sequencing Analysis Pipeline - GDC Docs. https://docs.gdc.cancer.gov/Data/Bioinformatics_Pipelines/DNA_Seq_Variant_Calling_Pipeline/.

36. Ellrott, K. et al. Scalable Open Science Approach for Mutation Calling of Tumor Exomes Using Multiple Genomic Pipelines. Cell Syst 6, 271–281.e7 (2018).

37. Bioinformatics Pipeline: Copy Number Variation Analysis - GDC Docs. https://docs.gdc.cancer.gov/Data/Bioinformatics_Pipelines/CNV_Pipeline/.

38. Mermel, C. H. et al. GISTIC2.0 facilitates sensitive and confident localization of the targets of focal somatic copy-number alteration in human cancers. Genome Biol 12, R41 (2011).

39. Bibikova, M. et al. High density DNA methylation array with single CpG site resolution. Genomics 98, 288–295 (2011).

40. Bioinformatics Pipeline: mRNA Analysis - GDC Docs. https://docs.gdc.cancer.gov/Data/Bioinformatics_Pipelines/Expression_mRNA_Pipeline/.

41. Dobin, A. et al. STAR: ultrafast universal RNA-seq aligner. Bioinformatics 29, 15–21 (2013).

42. Bioinformatics Pipeline: Protein Expression - GDC Docs. https://docs.gdc.cancer.gov/Data/Bioinformatics_Pipelines/RPPA_intro/.

43. Li, J. et al. TCPA: a resource for cancer functional proteomics data. Nature Methods 10, 1046–1047 (2013).

44. Akbani, R. et al. A pan-cancer proteomic perspective on The Cancer Genome Atlas. Nature Communications 5, 3887 (2014).

45. Kefeli, J. TCGA-Reports: A Machine-Readable Pathology Report Resource for Benchmarking Text-Based AI Models. Kefeli, et al. Mendeley Data 10.17632/HYG5XKZNPX.1 (2024).

46. Kefeli, J. & Tatonetti, N. TCGA-Reports: A machine-readable pathology report resource for benchmarking text-based AI models. Patterns 5, 100933 (2024).

47. Introducing Sonnet 4.6. https://www.anthropic.com/news/claude-sonnet-4-6.

48. Barbie, D. A. et al. Systematic RNA interference reveals that oncogenic KRAS-driven cancers require TBK1. Nature 462, 108–112 (2009).

