## Supplementary material for "Evaluating agentic AI for biological discovery in autonomous and copilot settings": Supplementary Note 1.pdf

### Appendix A

**AA3 Cell-Type Ontologies (per cohort)**

Annotations next to each node indicate which raw labels resolve to it:

  [AI: "..."]   raw AI vocabulary label

  [GT: "..."]   raw HTAN ground-truth label

Nodes without annotations are structural groupings used for hierarchy-aware grading.

#### BRCA

```
cell
├── epithelial
│   ├── epithelial_normal
│   │   ├── basal_progenitor  [GT: "Basal progenitor"]
│   │   ├── luminal_progenitor  [GT: "Luminal progenitor"]
│   │   └── luminal_mature  [GT: "Luminal mature"]
│   └── epithelial_tumor  [AI: "Cancer Epithelial" | GT: "Tumor"]
├── stromal
│   ├── fibroblast  [AI: "Fibroblast" | GT: "Fibroblasts"]
│   └── pericyte  [AI: "Pericyte" | GT: "Pericytes"]
├── endothelial  [AI: "Endothelial" | GT: "Endothelial"]
└── immune
    ├── lymphoid
    │   ├── b_cell  [AI: "B cell" | GT: "B-cells"]
    │   ├── t_cell  [AI: "T cell" | GT: "T-cells"]
    │   └── plasma_cell  [AI: "Plasma cell" | GT: "Plasma"]
    └── myeloid  [AI: "Myeloid"]
        ├── macrophage  [GT: "Macrophages"]
        ├── dendritic_cell  [AI: "Dendritic cell" | GT: "DC"]
        └── mast_cell  [AI: "Mast cell" | GT: "Mast"]
```

#### CESC

```
cell
├── epithelial  [AI: "Epithelial cell"]
│   ├── epithelial_normal
│   │   ├── normal_epithelial  [GT: "Normal epithelial cells"]
│   │   └── squamous  [GT: "Normal squamous cells"]
│   └── epithelial_tumor  [GT: "Tumor"]
├── stromal
│   └── fibroblast  [AI: "Fibroblast" | GT: "Fibroblasts"]
├── endothelial  [AI: "Endothelial cell" | GT: "Endothelial"]
└── immune
    ├── lymphoid
    │   ├── t_nk_cell  [AI: "T/NK cell"]
    │   │   └── t_cell  [GT: "T-cells"]
    │   ├── b_cell  [GT: "B-cells"]
    │   └── plasma_cell  [AI: "Plasma cell" | GT: "Plasma"]
    └── macrophage  [AI: "Macrophage" | GT: "Macrophages"]
```

CRC

```
cell
├── epithelial
│   ├── epithelial_normal
│   │   ├── intestinal
│   │   │   ├── distal_colon
│   │   │   │   ├── distal_enterocyte  [GT: "Distal Enterocytes"]
│   │   │   │   ├── distal_absorptive  [GT: "Distal Absorptive"]
│   │   │   │   ├── distal_ta  [GT: "Distal TA"]
│   │   │   │   ├── distal_mature_enterocyte  [GT: "Distal Mature Enterocytes"]
│   │   │   │   └── distal_stem  [GT: "Distal Stem Cells"]
│   │   │   ├── goblet  [GT: "Goblet"]
│   │   │   ├── ta  [GT: "TA"]
│   │   │   └── absorptive_best4_otop2  [GT: "Absorptive BEST4/OTOP2 cells"]
│   │   ├── liver
│   │   │   ├── cholangiocyte  [AI: "Cholangiocytes" | GT: "Cholangiocytes"]
│   │   │   └── hepatocyte  [AI: "Hepatocytes" | GT: "Hepatocytes"]
│   │   ├── alveolar  [GT: "Alveolar"]
│   │   └── mesothelial  [AI: "Mesothelial cells"]
│   └── epithelial_tumor  [AI: "Tumor cells" | GT: "Tumor"]
├── stromal
│   └── fibroblast  [AI: "Fibroblasts" | GT: "Fibroblasts"]
├── endothelial  [AI: "Endothelial cells" | GT: "Endothelial"]
└── immune
    ├── lymphoid
    │   ├── t_cell  [GT: "T-cells"]
    │   │   ├── cd4_t_cell  [AI: "CD4 T cells"]
    │   │   └── regulatory_t_cell  [AI: "Regulatory T cells"]
    │   ├── b_cell  [AI: "B cells" | GT: "B-cells"]
    │   └── plasma_cell  [AI: "Plasma cells" | GT: "Plasma"]
    └── macrophage  [AI: "Macrophages" | GT: "Macrophages"]
```

GBM

```
cell
├── neural
│   ├── glia
│   │   ├── astrocyte  [AI: "Astrocyte" | GT: "Astrocytes"]
│   │   ├── oligodendrocyte  [AI: "Oligodendrocyte" | GT: "Oligodendrocytes"]
│   │   └── opc  [AI: "OPC" | GT: "OPC"]
│   ├── neuron  [GT: "Neurons"]
│   │   ├── excitatory_neuron  [AI: "Excitatory Neuron"]
│   │   └── inhibitory_neuron  [AI: "Inhibitory Neuron"]
│   └── microglia  [AI: "Microglia" | GT: "Microglia"]
├── stromal
│   └── pericyte  [GT: "Pericytes"]
├── endothelial  [AI: "Endothelial"]
└── immune
    ├── lymphoid
    │   └── t_cell  [GT: "T-cells"]
    └── myeloid
        └── macrophage  [GT: "Macrophages"]
```

#### HNSCC

```
cell
├── epithelial  [AI: "Epithelial"]
│   ├── epithelial_normal  [AI: "Cycling Epithelial"]
│   └── epithelial_tumor  [GT: "Tumor"]
├── stromal
│   ├── fibroblast  [AI: "Fibroblast" | GT: "Fibroblasts"]
│   ├── pericyte  [AI: "Pericyte" | GT: "Pericytes"]
│   └── skeletal_muscle  [GT: "Skeletal Muscle"]
├── endothelial  [AI: "Endothelial" | GT: "Endothelial"]
└── immune
    ├── lymphoid
    │   ├── t_cell  [AI: "T cell" | GT: "T-cells"]
    │   ├── b_cell  [AI: "B cell" | GT: "B-cells"]
    │   └── plasma_cell  [AI: "Plasma cell" | GT: "Plasma"]
    └── myeloid
        ├── macrophage  [AI: "Macrophage" | GT: "Macrophages"]
        └── dendritic_cell  [GT: "DC"]
```

## OV

```
cell
├── epithelial  [AI: "Epithelial cell"]
│   └── epithelial_tumor  [GT: "Tumor"]
├── stromal
│   ├── fibroblast  [AI: "Fibroblast" | GT: "Fibroblasts"]
│   ├── pericyte  [AI: "Pericyte"]
│   └── adipocyte  [AI: "Adipocyte"]
├── endothelial  [AI: "Endothelial cell" | GT: "Endothelial"]
└── immune  [GT: "Immune"]
    ├── lymphoid
    │   ├── t_cell  [AI: "T cell" | GT: "T-cells"]
    │   ├── b_cell  [AI: "B cell" | GT: "B-cells"]
    │   └── plasma_cell  [AI: "Plasma cell" | GT: "Plasma"]
    └── macrophage  [AI: "Macrophage" | GT: "Macrophages"]
```

PDAC

```
cell
├── epithelial
│   ├── epithelial_normal
│   │   ├── pancreatic
│   │   │   ├── ductal  [AI: "Ductal"]
│   │   │   │   ├── ductal_like_1  [GT: "Ductal-like1"]
│   │   │   │   └── ductal_like_2  [GT: "Ductal-like2"]
│   │   │   ├── acinar  [AI: "Acinar" | GT: "Acinar"]
│   │   │   │   └── acinar_reg_pos  [GT: "Acinar REG+"]
│   │   │   └── endocrine  [AI: "Endocrine"]
│   │   │       └── islet  [GT: "Islets"]
│   │   └── hepatocyte  [AI: "Hepatocyte" | GT: "Hepatocytes"]
│   └── epithelial_tumor
│       ├── tumor  [GT: "Tumor"]
│       └── adm  [GT: "ADM"]
├── stromal
│   ├── fibroblast  [AI: "Fibroblast" | GT: "Fibroblasts"]
│   └── adipocyte  [GT: "Adipocytes"]
├── endothelial  [AI: "Endothelial" | GT: "Endothelial"]
├── cycling  [AI: "Cycling"]
└── immune
    ├── lymphoid
    │   ├── t_cell  [AI: "T cell" | GT: "T-cells"]
    │   ├── b_cell  [AI: "B cell" | GT: "B-cells"]
    │   └── plasma_cell  [AI: "Plasma" | GT: "Plasma"]
    ├── macrophage  [AI: "Macrophage" | GT: "Macrophages"]
    └── mast_cell  [AI: "Mast cell" | GT: "Mast"]
```

#### SKCM

```
cell
├── epithelial
│   ├── epithelial_normal
│   │   ├── keratinocyte  [GT: "Keratinocytes"]
│   │   └── melanocyte  [GT: "Melanocytes"]
│   └── epithelial_tumor  [AI: "Melanoma cell" | GT: "Tumor"]
├── stromal  [AI: "Stromal cell"]
│   ├── fibroblast  [AI: "Fibroblast" | GT: "Fibroblasts"]
│   │   └── myofibroblast  [AI: "Myofibroblast"]
│   ├── pericyte  [AI: "Pericyte"]
│   └── smooth_muscle  [GT: "Smooth muscle"]
│       └── vsmc  [GT: "vSMCs"]
├── endothelial  [AI: "Endothelial cell" | GT: "Endothelial"]
└── immune
    ├── lymphoid
    │   ├── t_cell  [AI: "T cell" | GT: "T-cells"]
    │   ├── b_cell  [AI: "B cell" | GT: "B-cells"]
    │   └── plasma_cell  [AI: "Plasma cell" | GT: "Plasma"]
    ├── myeloid
    │   ├── macrophage  [AI: "Macrophage" | GT: "Macrophages"]
    │   └── dendritic_cell  [AI: "Dendritic cell" | GT: "DC"]
    │       └── pdc  [AI: "pDC"]
    └── mast_cell  [GT: "Mast"]
```

#### UCEC

```
cell
├── epithelial  [AI: "Epithelial cell"]
│   ├── epithelial_normal  [AI: "Cycling epithelial cell"]
│   │   ├── secretory_endometrial  [GT: "Secretory Endometrial epithelial cells"]
│   │   └── ciliated_endometrial  [AI: "Ciliated epithelial cell" | GT: "Ciliated Endometrial epithelial cells"]
│   └── epithelial_tumor  [GT: "Tumor"]
├── stromal
│   ├── fibroblast  [AI: "Fibroblast" | GT: "Fibroblasts"]
│   └── pericyte  [AI: "Pericyte"]
├── endothelial  [AI: "Endothelial cell" | GT: "Endothelial"]
└── immune
    ├── lymphoid
    │   ├── t_cell  [AI: "T cell" | GT: "T-cells"]
    │   ├── b_cell  [AI: "B cell" | GT: "B-cells"]
    │   └── plasma_cell  [AI: "Plasma cell" | GT: "Plasma"]
    ├── myeloid
    │   ├── macrophage  [AI: "Macrophage" | GT: "Macrophages"]
    │   └── dendritic_cell  [GT: "DC"]
    └── mast_cell  [GT: "Mast"]
```
