## Supplementary material for "Evaluating agentic AI for biological discovery in autonomous and copilot settings": Supplementary_Figures.pdf

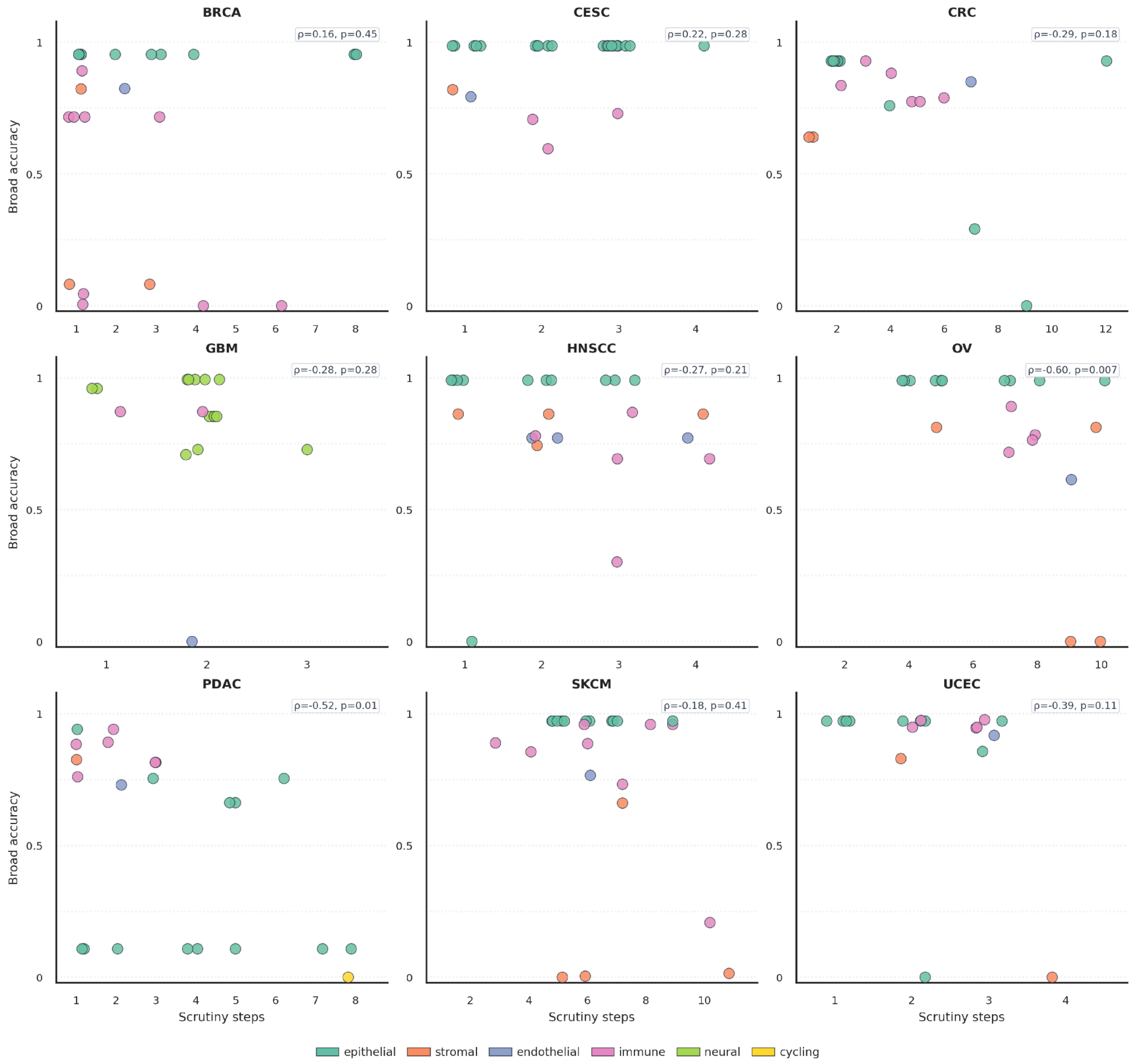


**Supplementary Figure 1:** Per-cluster scrutiny depth versus broad annotation accuracy, stratified by cancer type (BRCA, CESC, CRC, GBM, HNSCC, OV, PDAC, SKCM, UCEC) and cell-type lineage (epithelial, stromal, endothelial, immune, neural, cycling).

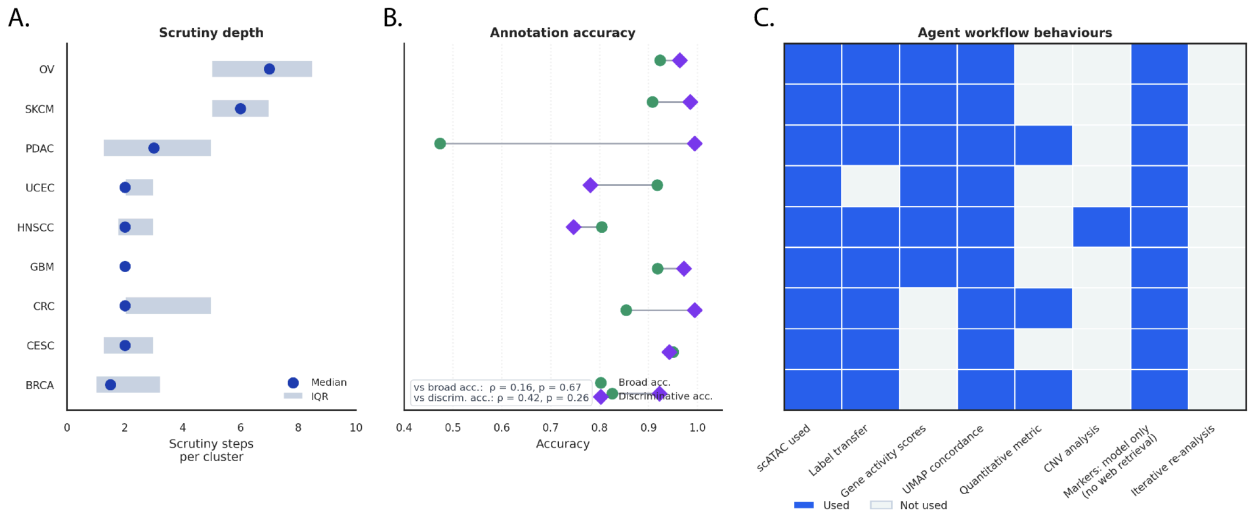


**Supplementary Figure 2:** M3A agent scrutiny depth, annotation accuracy, and workflow strategy (scATAC usage, label transfer, gene activity scoring, UMAP concordance, quantitative metrics, copy-number variation (CNV) analysis, knowledge driven marker analysis, iterative re-analysis) by cancer type (OV, SKCM, PDAC, UCEC, HNSCC, GBM, CRC, CESC, BRCA).

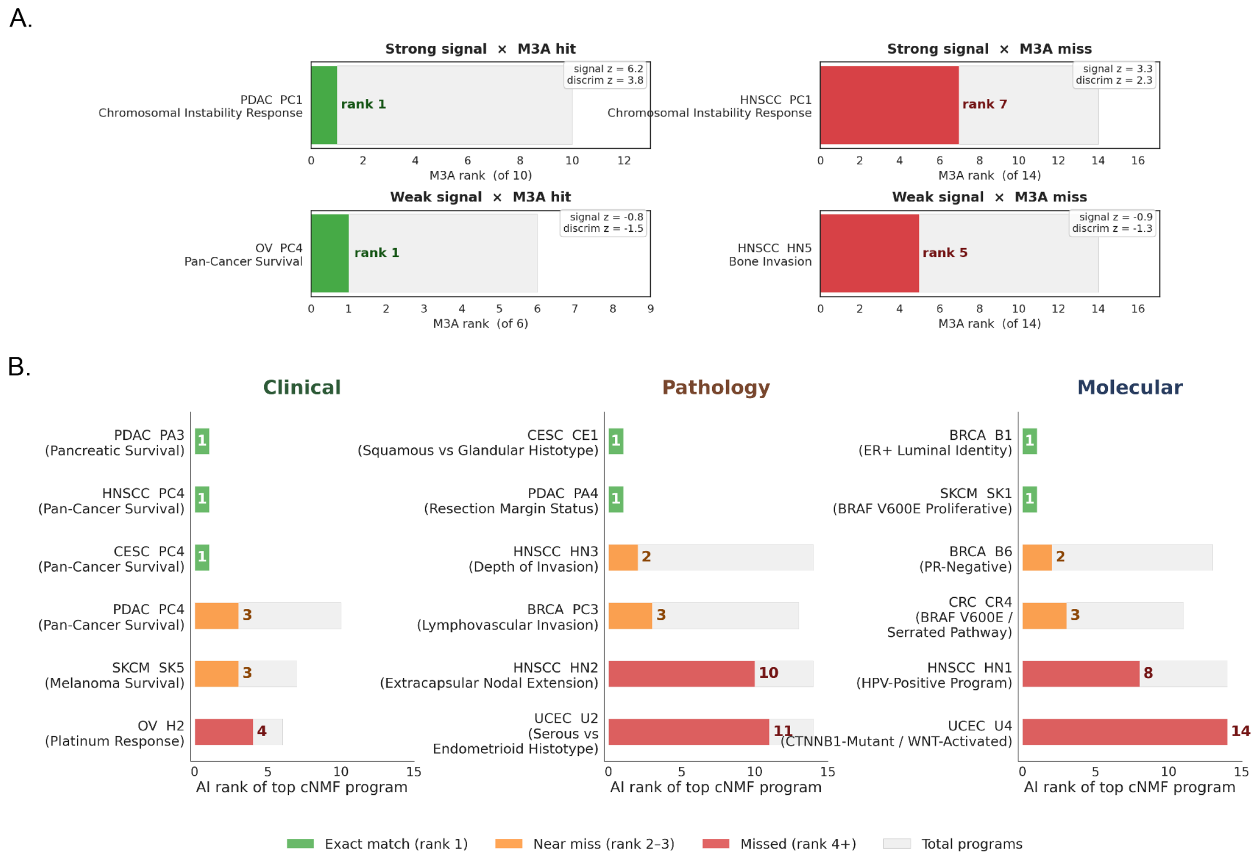


**Supplementary Figure 3: (a)** Representative tasks from each quadrant of the signal-strength × M3A-agent’s outcome space (M3A hit, M3A miss); light-gray bar indicates the total number of cNMF programs in that task. **(b)** Representative example tasks of M3A-agent prioritization being an exact match (green, rank 1), near miss (orange, rank 2-3), or clear miss (red, rank ≥ 4) of the statistically top-1 program; light-gray bar indicates the total number of cNMF programs in that task.

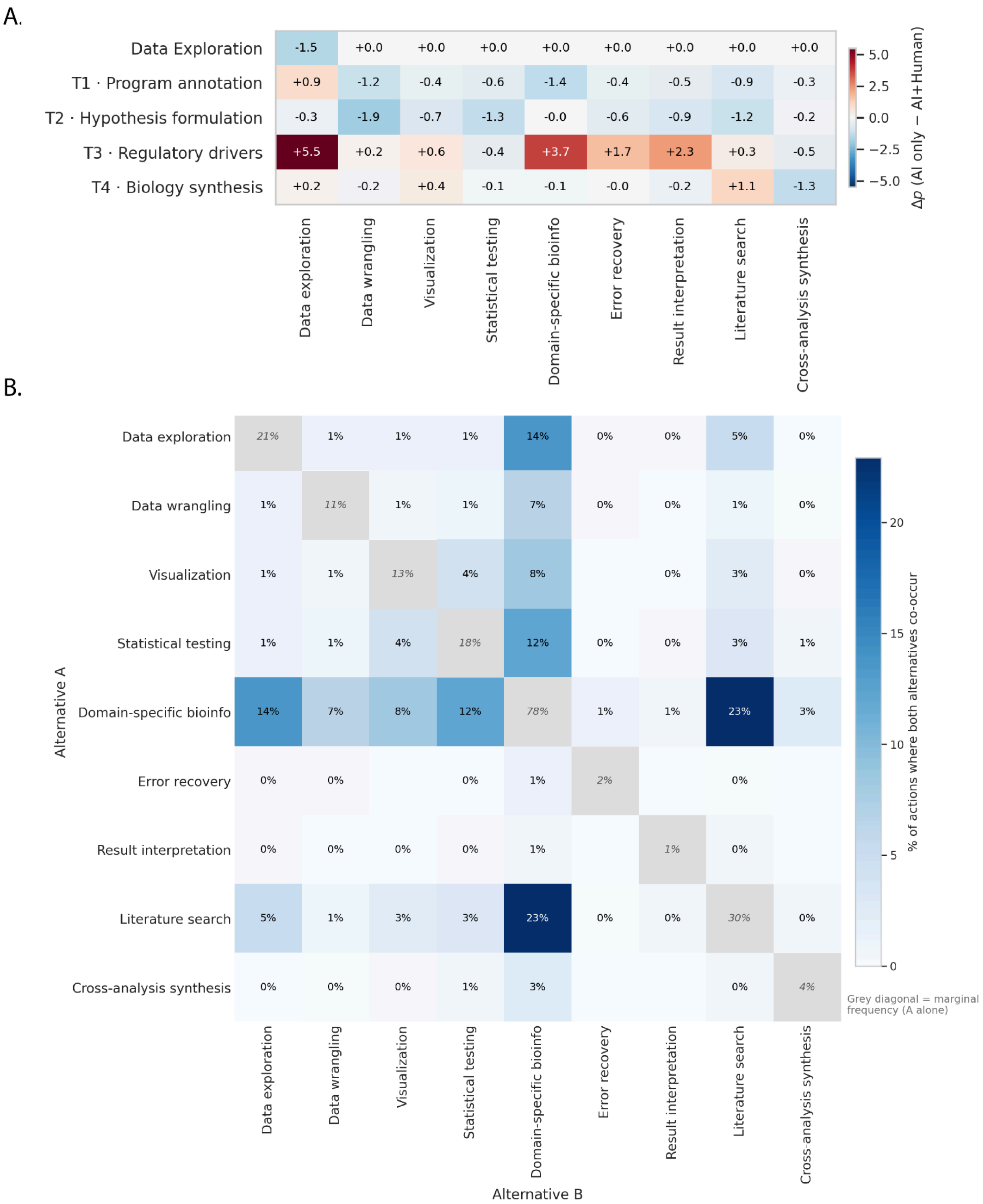

**Supplementary Figure 4: (a)** M3A agent action category (data exploration, data wrangling, visualization, statistical testing, domain-specific bioinformatics, error recovery, result interpretation, literature search, cross-analysis synthesis) composition between AI-only and human+AI modes, stratified by tasks (Data exploration; Program annotation, T1; Hypothesis formulation, T2; Regulatory driver analysis, T3; Biology synthesis, T4). **(b)** Co-occurrence of alternative M3A agent actions.

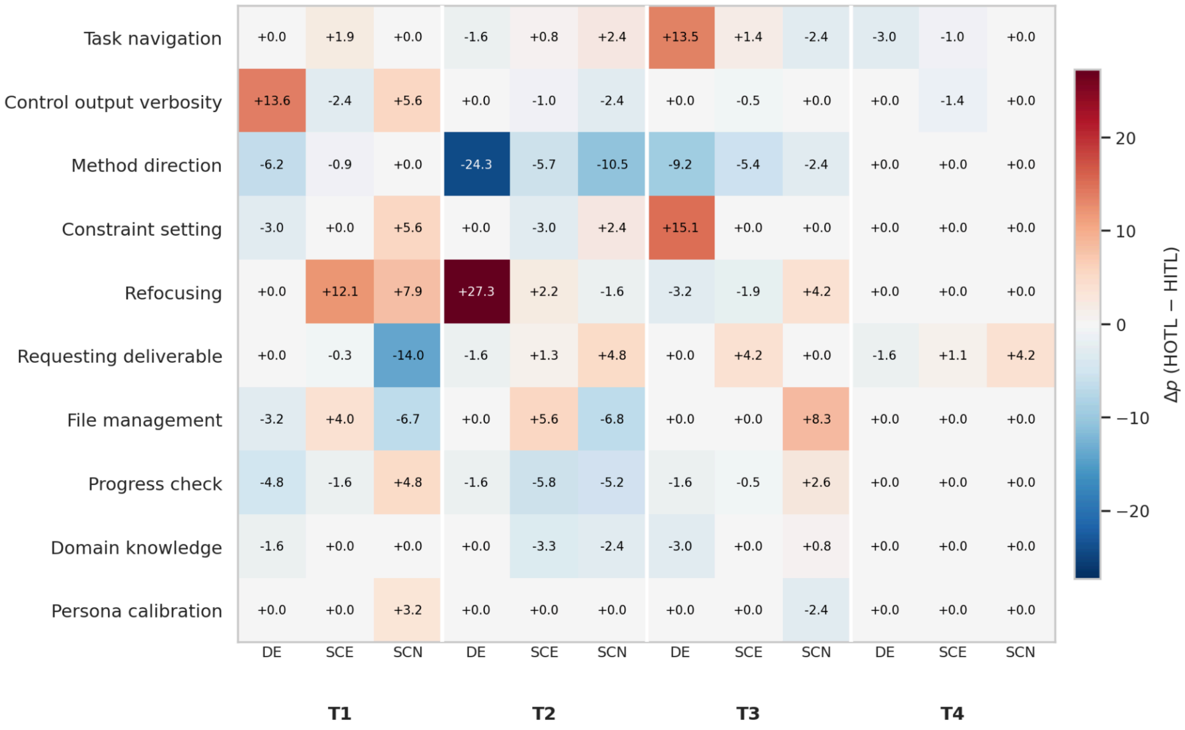


**Supplementary Figure 5:** Impact of interaction mode (HOTL - HITL) in user interaction category (task navigation, output formatting, method direction, constraint setting, correcting/refocusing, requesting deliverable, file/output management, progress check, domain knowledge, personal calibration) composition, stratified by task stage (Program annotation, T1; Hypothesis formulation, T2; Regulatory driver analysis, T3; Biology synthesis, T4) and experience level (Domain Expert (DE), SC Experienced(SCE), SC Novice(SCN)).

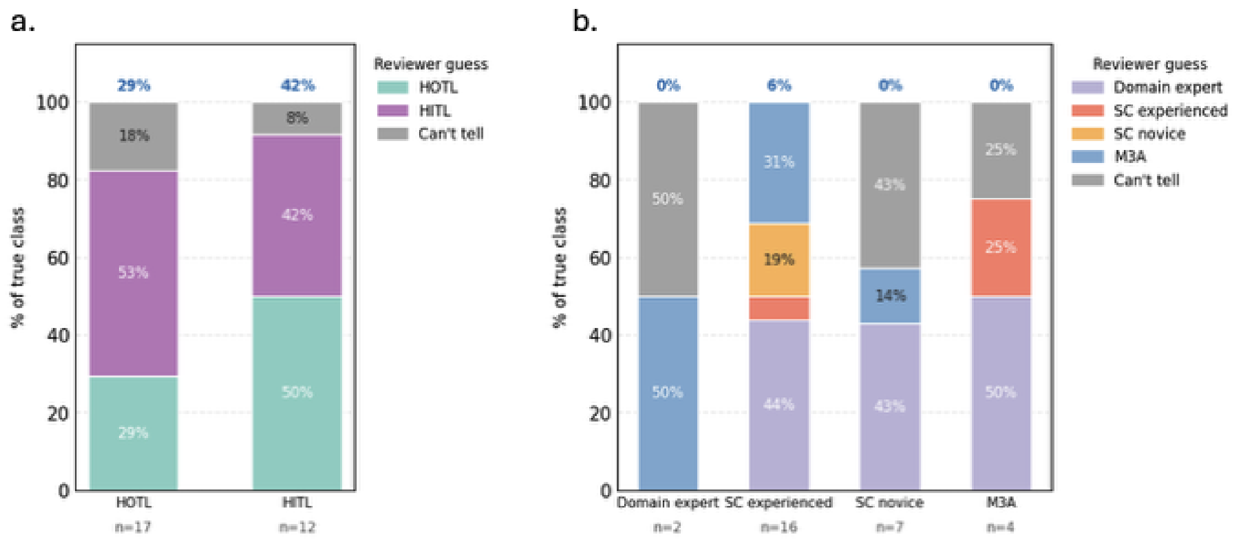


**Supplementary Figure 6:** Reviewer identification of **(a)** interaction mode and **(b)** analyst background. Each bar represents a true interaction style (HOTL or HITL) or a true analyst background category (Domain expert, SC experienced, SC novice, M3A) and shows the distribution of reviewer guesses across all cancer cohorts pooled.
